# The theory and practice of measuring broad-range recombination rate from marker selected pools

**DOI:** 10.1101/762575

**Authors:** Kevin H.-C. Wei, Aditya Mantha, Doris Bachtrog

## Abstract

Recombination is the exchange of genetic material between homologous chromosomes via physical crossovers. Pioneered by T. H. Morgan and A. Sturtevant over a century ago, methods to estimate recombination rate and genetic distance require scoring large number of recombinant individuals between molecular or visible markers. While high throughput sequencing methods have allowed for genome wide crossover detection producing high resolution maps, such methods rely on large number of recombinants individually sequenced and are therefore difficult to scale. Here, we present a simple and scalable method to infer near chromosome-wide recombination rate from marker selected pools and the corresponding analytical software MarSuPial. Rather than genotyping individuals from recombinant backcrosses, we bulk sequence marker selected pools to infer the allele frequency decay around the selected locus; since the number of recombinant individuals increases proportionally to the genetic distance from the selected locus, the allele frequency across the chromosome can be used to estimate the genetic distance and recombination rate. We mathematically demonstrate the relationship between allele frequency attenuation, recombinant fraction, genetic distance, and recombination rate in marker selected pools. Based on available chromosome-wide recombination rate models of *Drosophila*, we simulated read counts and determined that nonlinear local regressions (LOESS) produce robust estimates despite the high noise inherent to sequencing data. To empirically validate this approach, we show that (single) marker selected pools closely recapitulate genetic distances inferred from scoring recombinants between double markers. We theoretically determine how secondary loci with viability impacts can modulate the allele frequency decay and how to account for such effects directly from the data. We generated the recombinant map of three wild derived strains which strongly correlates with previous genome-wide measurements. Interestingly, amidst extensive recombination rate variation, multiple regions of the genomes show elevated rates across all strains. Lastly, we apply this method to estimate chromosome-wide crossover interference. Altogether, we find that marker selected pools is a simple and cost effective method for broad recombination rate estimates. Although it does not identify instances of crossovers, it can generate near chromosome-wide recombination maps in as little as one or two libraries.

## INTRODUCTION

T. H. Morgan first envisioned the exchange of genetic material between homologous chromosomes through physical crossovers (Morgan 1911b). After backcrossing F1 heterozygotes, he recovered novel allelic combinations, or recombinants, of markers on the same chromosome that are absent in parental lines (Morgan 1911a), a violation, at face value, to Mendal’s law of segregation and the chromosome theory of inheritance (Blixt 1975). A. H. Sturtevant, then a prodigious undergraduate in Morgan’s laboratory, confirmed his mentor’s theory by crossing mutant *Drosophila* strains with different visible X-linked markers. Noticing that different pairs of markers produced recombinants at different frequencies, Sturtevant realized that the positions of these markers can be ordered on a linear map, with the frequency of recombinants as the genetic distance between any two markers (Sturtevant 1913). This was the birth of the very first genetic map.

While this might seem incontrovertible in hindsight, Morgan and Sturtevant were not free of critics (Castle 1919; Brush 2002). One main point of contention was the fact that the frequency of recombinants (henceforth recombinant fraction) estimated as the number of recombinants divided by the total number of offsprings is nonadditive. Given three loci, A, B, and C in that order, the recombinant fraction between A and C is typically shorter than the sum of A-to-B and B-to-C. As pointed out by Sturtevant, Bridges and Morgan, the reason for this is that between any two points, the recombinant fraction only captures crossover that generate visible recombinants; as such, even numbers of crossovers are unobservable, and as genetic distances increase (as in A-to-C), the chances of even numbers of crossovers also increase, resulting in the nonadditivity (Sturtevant *et al.* 1919). J. B. S. Haldane formally described the relationship between true genetic distance (*d*), which is the frequency of crossovers between two loci, and recombinant fraction (*D*), which is the frequency of odd number of (observable) crossovers, with the formulae:

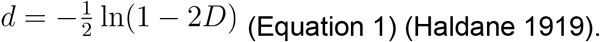

After the transformation with Haldane’s mapping function (**H**) non-additive recombinant fraction than become additive genetic distance such that:

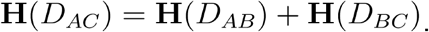

According to this function, the recombinant fraction and genetic distance are near identical at low values (< 0.1) reflecting the fact that even-numbered crossovers have negligible frequencies, but as the former increases, the latter increases exponentially, approaching infinity at recombinant fraction of 0.5. Haldane’s mapping function is effectively capturing the nonadditive property of recombinant fraction with the following:

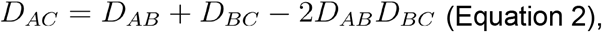

where the recombinant fraction between A and C is the sum of the recombinant fraction between the two constituent fractions minus the probability of double crossovers. Thus, Haldane’s function assumes that crossovers are independent of each other, but unbeknownst to him at the time, the probabilities crossovers are not independent.

Based on extensive three point crosses experiments (i.e. crosses with three visible markers) in many organisms, it became clear that the fraction of double crossovers is typically less than the expectation (*2D*_*AB*_*D*_*BC*_), revealing that once a single crossover occurs, a second crossover is less likely (Sturtevant 1913; Muller 1916; Baker *et al.* 1976; Hawley 1988; Hillers 2004). This phenomenon, known as crossover interference, led to the adjustment of the non-additive relationship to:

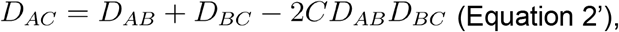

where *C*, known as the coefficient of coincidence and is typically between 0 and 1, modifies the rate of double crossovers; crossover interference is then 1 − *C*. When *C* is 0, i.e. full interference, the recombinant fraction is fully additive and equals the genetic distance, but when *C* is 1, i.e. no interference, the Haldane mapping function is true. Without accounting for interference, Haldane’s mapping function results in overestimated genetic distances particularly with larger recombinant fractions (Tan and Fornage 2008).

Incorporation of interference into the mapping function came decades later from the mathematician D. D. Kosambi (Kosambi 1943). He sensibly assumed that crossover interference is proportional to the recombinant fraction with the second crossover probability fully recovered at a distance of 0.5, resulting in the simple relationship of *C* = *2D* and non-additivity of:

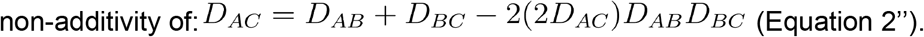

Incorporating this, he derived the new mapping function (**K**):

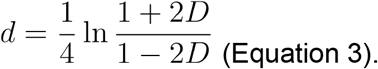

Compared to Haldane’s function, Kosambi’s function, similarly, provides negligible changes at low recombinant fractions, but produces shorter genetic distances at higher recombinant fractions and are more consistent with results from crossing experiments (Huehn 2011). While other mapping functions have been developed to account for different extent of interference (Felsenstein 1979; McPeek and Speed 1995; Tan and Fornage 2008), Kosambi’s and Haldane’s functions remain among the most popular.

In the century after Morgan and Sturtevant conceptualized and devised the recombinant backcross scheme, recombination has been recognized to be one of the most fundamental and universal biological processes across sexually reproducing eukaryotes with vast implications in many biological fields. For example, proper assembly of the complex crossover machinery is crucial for the fidelity of chromosome segregation (see reviews (Page and Hawley 2004), (Hunter 2015), and (Hughes *et al.* 2018)) and recombination rate is intimately linked with the efficacy of natural selection and genome evolution (see reviews (Cutter and Payseur 2013), (Martin and Jiggins 2017), and (Stephan 2019)). Despite rapid technological and methodological advances in these relevant fields, methods to measure recombinant fraction, genetic distance, and recombination rate remain grounded by the approach devised by Sturtevant - tallying the number of recombinants between loci that are either phenotypically or molecularly marked. The advent of whole genome sequencing has permitted high resolution crossover maps, by identifying regions of the genome where parental haplotypes change in recombinant individuals (Kulathinal *et al.* 2008; Rockman and Kruglyak 2009; Dumont *et al.* 2011; Miller *et al.* 2012; Comeron *et al.* 2012). More recently, single-cell sequencing technologies have further extended such high-throughput crossover detection directly from sperms (Hinch *et al.* 2019). While providing impressive resolution often at the level of individual bases, these approaches are difficult to scale, as each recombinant individual/cell requires a separate library preparation and/or barcode. Since each chromosome, on average, has on average one to two crossovers, large numbers of individuals, and therefore libraries/barcodes, need to be sequenced for a comprehensive map. Given the environmentally-induced volatility (Stern 1926; Neel 1941; Redfield 1966) and inter-(Jensen-Seaman *et al.* 2004; Kulathinal *et al.* 2008; Smukowski and Noor 2011; Brand *et al.* 2018) and intra-specific variability (Nachman 2002; Dumont *et al.* 2009, 2011; Comeron *et al.* 2012; Kaur and Rockman 2014) of recombination rates, a scalable high-throughput method of estimation will greatly facilitate our characterization and understanding of recombination.

Here, we present a simple marker selection followed by pooled sequencing scheme to estimate chromosome wide recombinant fraction, from which genetic distance and recombination rate can be inferred. The key benefit of this approach is that it requires orders of magnitude fewer libraries/barcodes than previous approaches. In a typical recombinant backcross scheme to estimate genetic distance between two markers, the allele frequency (AF) of the entire progeny pool is expected to be the same across the chromosome, despite the presence of recombinants that resulted from crossovers in the F1 parents (Figure 1A). However, if the progeny are sorted into separate pools based on a marker, the AF is expected show a signature pattern: the AF of the marked chromosome peaks at the marker locus and attenuates distally. This distal attenuation is the result of increasing number of crossover events between the marked locus and increasingly distal loci. Therefore, the allele frequency and its rate of change should be related to the genetic distance and recombination rate, respectively. Previously, allele frequency has also been utilized for fine scale crossover rate estimates in pools where recombinants were specifically selected for (Singh *et al.* 2013) but had analytical shortfalls that resulted in problematic estimates (Gilliland 2015) (also see Discussion). Our approach of marker selection does not require any identification of recombinants.

**Figure 1.**
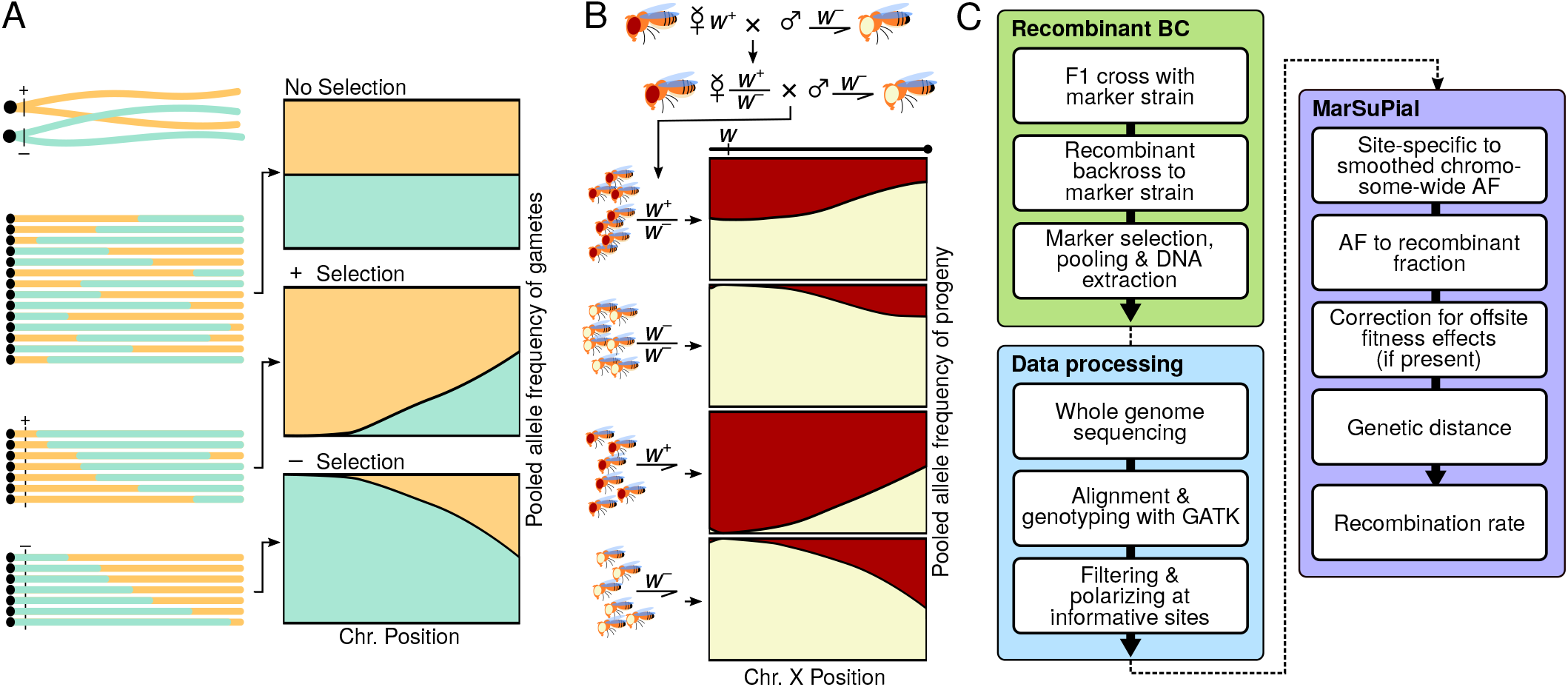
Marker selection pooling scheme to estimate recombination rate. A. Rationale of the marker selection scheme to create allele frequency attenuation. On the left are cartoon schematics of crossover between homologous chromosomes and their recombinant products. After marker selection, the recombinant chromosomes are ordered by the length of the haplotypes for illustrative purposes. The allele frequency of the recombinant pools are depicted in the cartoons to the right. B. Application of this approach using the X-linked eye color marker *white* (*w*). The recombinant backcross schemes is on the top. The resulting four sexed and genotyped BC1 pools are displayed below; the allele frequency of these pools are displayed to the right, where the red and white areas represent the frequency of the *w*^+^ and *w^−^* alleles, respectively. C. Workflow of the method, from fly crosses to recombination rate estimation using the Software package MarSuPial.

We theoretically and empirically explore the series of steps to infer recombination rates from allele frequency attenuation in marker selected pools. First, we formally demonstrate the relationship between recombination rate and allele frequency in the context of the Drosophila X chromosome with the recessive white eye marker (w^−^). We show how inherent noise in the sequencing platform can be addressed statistically. We then empirically show that genetic distances estimated from the allele frequency changes closely recapitulate the distances estimated from the de facto approach of scoring recombinant individuals, validating the efficacy of this method. Anticipating fitness impacts from offsite loci, we demonstrate theoretically how they modulate the allele frequency and how they can be corrected from data. Using this approach, we generated the 3rd chromosome recombination map of three wild derived strains, producing rates that are highly correlated with previous estimates. Lastly, we extend the application of this approach to estimate crossover interference across the 3rd chromosome. While this marker selection and sequencing scheme does not produce high-resolution maps since it does not infer breakpoints, it can generate a near chromosome-wide genetic map (particularly in *D. melanogaster*) in as little as one or two libraries.

## RESULTS

### Genetic distance is directly proportional to recombinant fraction in marker selected pool

While gametes are ideal for detecting crossovers as they are direct products of meiosis (Figure 1A), there is currently no viable way of gamete selection. In practice, marker selection is accomplished in the backcross offsprings of the F1 mothers and half of the offspring genomes originate from the inbred father and is uninformative, despite contributing to the allele frequency. However, since the X chromosome in sons are maternally inherited, male and female pools will have different allele frequency signatures (Figure 1B). For simplicity, we will first focus on the male pools as their X chromosomes are free of paternal contributions and therefore represents the gametic AF. For illustrative purposes, we will use the X-linked white gene in *Drosophila* as the selected locus (*w*) (Figure 1B); selection is based on the recessive phenotype of white eyes (*w^−^*) versus red eyes (w^+^). Between the selected locus *w* and any position *i*, the recombinant frequency between the two loci is *D*_*i*_; the four possible allelic combinations will simply have the following frequencies (*p* and *q*, for the *w^+^* and *w^−^* chromosomes, respectively):

*p*_*i*_ = (1 − *D*_*i*_)/2 for the *w^+^* chromosomes with zero or even numbers of crossovers,

*q*_*i*_ = (1 − *D*_*i*_)/2 for the *w^−^* chromosomes with zero or even numbers of crossovers,

*p*_*i*_ = *D*_*i*_/2 for the *w^+^* chromosomes with odd numbers of crossovers.

*q*_*i*_ = *D*_*i*_/2 for the *w^−^* chromosomes with odd numbers of crossovers.

In the absence of selection, The AF of the *w^−^* chromosome chromosome-wide will therefore be:

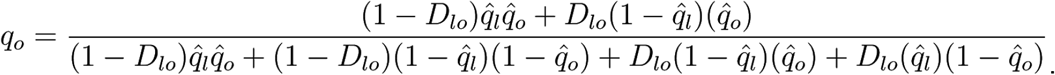

However, when *w^−^* is selected and pooled:

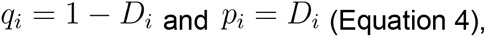

and when *w^+^* is selected and pooled:

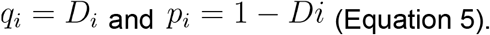

Therefore, the AF is directly proportional to the recombinant fraction. To illustrate this relationship between recombination and AF change in the male selected pools, we used the available X chromosome recombination rate from the Recombination Rate Calculator (Fiston-Lavier *et al.* 2010) which models the chromosome-wide rate by a quadratic function. We converted the recombination rate to genetic distance from the white locus, followed by transformation into the recombinant fraction using either the Haldane or Kosambi mapping functions (Figure 2A). Depending on whether *w^−^* and w+ males are selected, AF peaks (*q* = 1) or troughs (*q* = 0) at the selected white locus, respectively, and attenuates distally, asymptoting towards the expected Mendelian ratio of 0.5 (Figure 2B).

**Figure 2.**
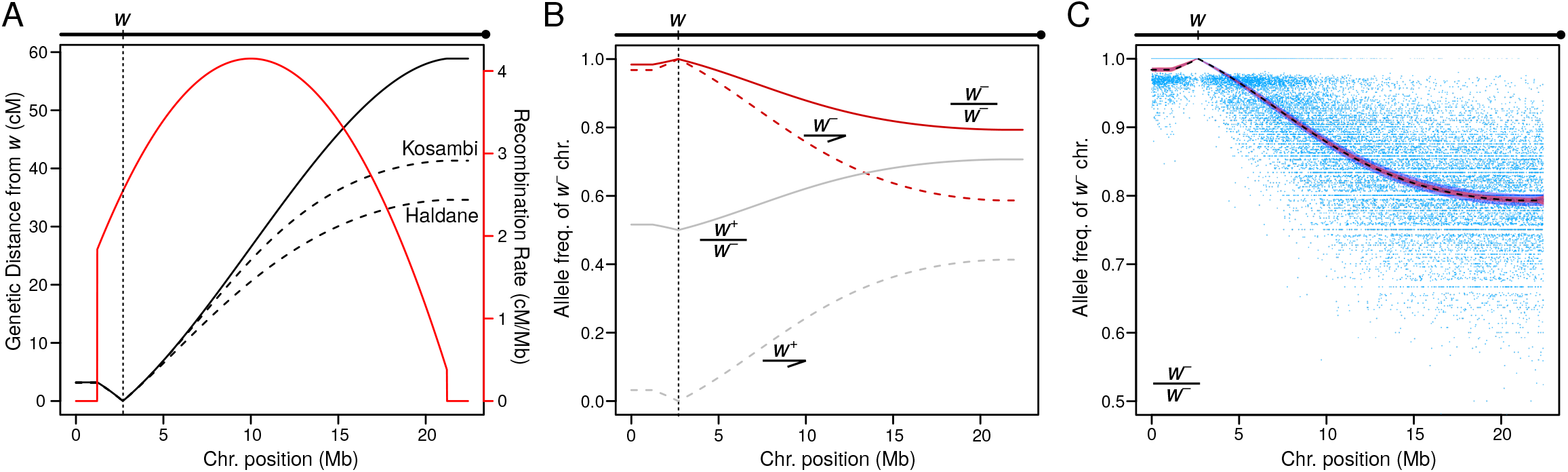
Mathematical evaluation and demonstration based on known recombination rates. A. Based on the backcross scheme in 1B, transformation between the recombination rate of the X chromosome (red line), the genetic distance from the white locus (solid black line), and the recombinant fractions after applying the Kosambi and Haldane mapping functions. B. Allele frequency of different pools across the X chromosome given the Kosambi transformed recombinant fraction from A. The *w^−^* and *w*^+^ pools are in gray and red lines, respectively; the female and male pools are in solid and dotted lines, respectively. C. Allele frequency from simulated read counts of the *w^−^* female pool. Given the SNP density of 1 site per 1000bp, the allele-specific read counts across the chromosome is randomly sampled to simulate fly collection, library preparation, and sequencing. 20000 trials were conducted to simulate 1000 pooled individuals and sequence depth of 30; the allele frequency at each site for one trial is plotted (blue) to illustrate the extent of noise from site to site. For each trial, the allele frequency chromosome-wide is either binned in overlapping 300kb windows followed by linear regression fit or smoothed with LOESS local regression anchored at the selected *w* locus. The true allele frequency (from 2B) is plotted in dotted black line. The 95% confidence interval of the linear regression and LOESS fits are demarcated by the blue and red areas around the AF, respectively.

In order to select for the recessive *w^−^* marker in females, they must be sired by homozygous *w^−^* fathers (Figure 1B). The *w^−^* chromosome allele frequency in the *w^−^* and *w^+^* pools with paternal contributions will then be:

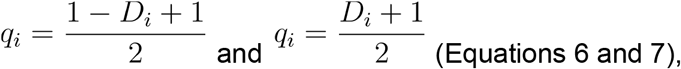

respectively, in females. The AF thus peaks (*q* = 1) or troughs (*q* = 0.5) at the selected white locus depending on whether homozygotes (*w^−^*/*w^−^*) or heterozygotes (*w^+^*/*w^−^*) are selected, respectively (Figure 2B) and attenuates toward the expected Mendelian ratio (*q* = 0.75).

To evaluate whether short read whole genome sequencing can sensitively capture the AF signature, we used Monte Carlo simulations varying the pool size and average sequencing depth with a conservative SNP density of 1 differentiating site per 1kb (see Materials and Methods). Within reasonable range of sequencing depth of 10x-100x, allele counts at individual sites are too variable to provide meaningful AF estimates (Figure 2C, Supplementary figure 1A). Since neighboring sites are expected to have negligible differences in their AFs, their counts and frequencies can be aggregated as if they are independent in sliding windows to better approximate the AF (Wei *et al.* 2017). However, instead of assuming all sites in a window have the same AF, we first estimated AF using linear regression in each overlapping sliding window (see Materials and Methods). We find that 500kb overlapping windows in 100kb intervals sensitively recapitulate the expected AF (Figure 2C, Supplementary figure 1B) but poorly estimates the slope of the AF decay, frequently producing the wrong sign (Supplementary figure 2). Lastly, rather than binning the genome into windows, we used local regressions (LOESS) (Cleveland 1979), one on either side of the selected locus, for non-parametric and non-linear curve fitting (Figure 2C, Supplementary figure 1B). We find the LOESS fit to be the most flexible and robust, as it allows for non-linear fit, is most robust to noise (Supplementary Figure 1B), and can accurately estimate the slope of the AF decay curve (Supplementary figure 2). Notedly, the variances of the estimated AF from both methods increase distally as AF tends toward Mendelian ratios; this is because variance is highest for binomial distributions with the intermediate probability of *p* = 0.5.

### Allele frequency from whole genome sequencing of marker selected pools closely estimates recombinant fraction

To empirically test the efficacy of this approach, we set up recombinant back crosses pairing two double recessive marker strains (sepia ebony (*se^−^ e^−^*) and ebony glass (*e^−^ gl^−^*)) with three wild-derived inbred strains (Canton-S, DGRP-315, and DGRP-360) (Table 1). We first sexed and tallied the number of recombinant and nonrecombinant individuals to infer the genetic distance between the two linked markers, then we pooled the flies based on the presence of one of the two markers followed by bulk DNA extraction and whole genome sequencing to an average of 25.45x coverage (Supplementary table 1). Thus, we were able to compare the genetic distance estimates between the two loci based on the de facto method of fly scoring to that from AF attenuation in the marker selected pool from the same recombinant backcrosses. Canton-S was separately crossed to the two double marker strains in duplicates; in the cross to *e^−^ gl^−^*, we selected and pooled *gl^−^* females but *e^−^* males with the expectation that the two pools should yield similar results. For DGRP-315 and DGRP-360, we crossed each of the two only to *e^−^ gl^−^* and pooled the *gl^−^* individuals. However, with the DGRP-315 cross, in addition to the *gl^−^* pool (positive marker selection) we also pooled the *gl*^+^ individuals (negative marker selection) from the same cross, again with the expectation that the two pools should yield identical results. Allele-specific read counts were determined only at informative SNP sites where the parental strains are homozygous for different nucleotides (see Materials and Methods) (Supplementary table 2 and Supplementary figure 3). Sites with segregating variants in the parental strains are removed (Supplementary table 3 and Supplementary figure 4). We then estimated the AF using LOESS fit as described above (also see Materials and Methods).

**Table 1.**
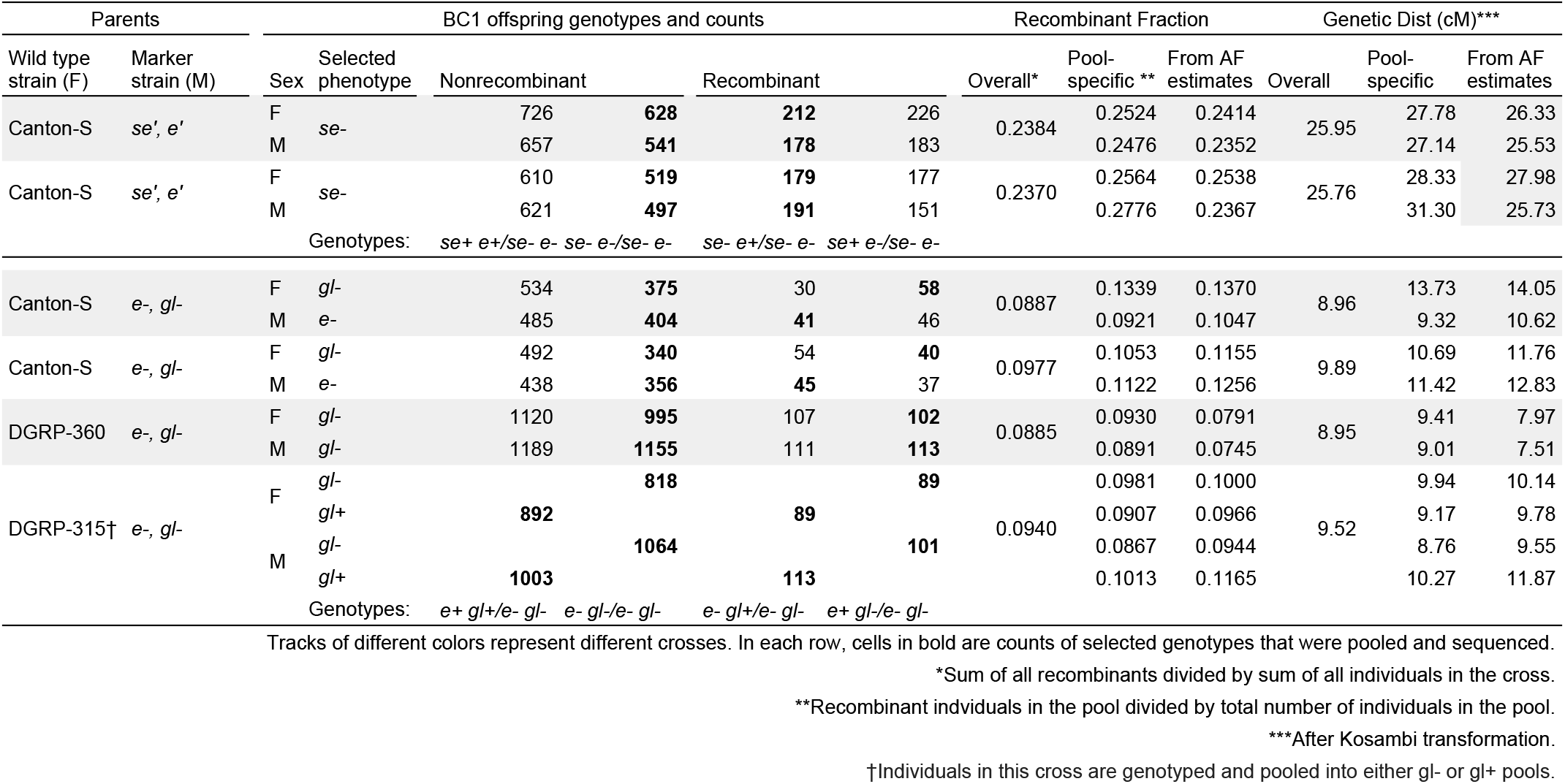
Genetic distance estimates between double markers

We find that the recombinant fractions and genetic distances determined from the two approaches are highly similar with no significant differences (p = 0.9515, Paired Wilcoxon Ranked Sum Test) (Table 1); with the exception of one pool (Canton-S x *se*^−^ *e*^−^), estimates are within 1.5% of each other which is comparable to the variability between fly count data of replicates as well as different pools from the same cross. Extensively, this demonstrates that the recombinant fraction can be inferred between the selected locus and any site along the chromosome until reaching 0.5 (Figure 3A-D, Supplementary figure 4). When individuals homozygous for the markers are pooled, the AFs of the chromosome carrying the marker peaks at the selected locus with AF of 1 and decreases distally (Figure 3A and C, Supplementary figure 3), consistent with the theoretical and simulated results. For DGRP315 x *gl^−^ e^−^* cross, the AF of the negative marker selection pool (*gl^−^*/*gl^+^*) expectedly dips to 0.5 at *gl* and increases distally (Figure 3D). In all these crosses, the AF stays at a near constant level across the centromere and pericentromeres of 3L and 3R, fitting the expectation of no recombination across the region. When converted to recombinant fractions, both the positive (*gl^−^*/*gl^−^*) and negative selection (*gl^−^*/*gl*^+^) pools yield highly similar results (Supplementary figure 5). Across replicates, and different pools from the same cross, the recombinant fraction appears to be the most variable around the pericentric region (Figure 3E-G), resulting from the reduced SNP density and poor mapping quality due to increased repeat content (Supplementary figure 6). However, the recombinant fractions converge outside of the pericentromere, producing highly robust estimates (Figure 3E-G). Notably, the Canton-S x *gl^−^ e^−^* crosses have the highest variance; this it due to the fact that the Canton-S strain used had unexpectedly high levels of heterozygosity which reduces the number of usable informative sites (Supplementary Table 3 and Supplementary figure 2 and 7). The genetic distance can then be estimated by transformation with mapping functions (Figure 3E-G, dotted lines). Necessitated by these functions, while the recombinant fraction approaches 0.5, the genetic distance approaches infinity. There, the genetic distance should be, in practice, limited to 50cM which entails that two loci are effectively genetically unlinked (Figure 3E-G); this equates to recombinant fractions of < 0.381 and 0.316 with the Kosambi and Haldane functions, respectively.

**Figure 3.**
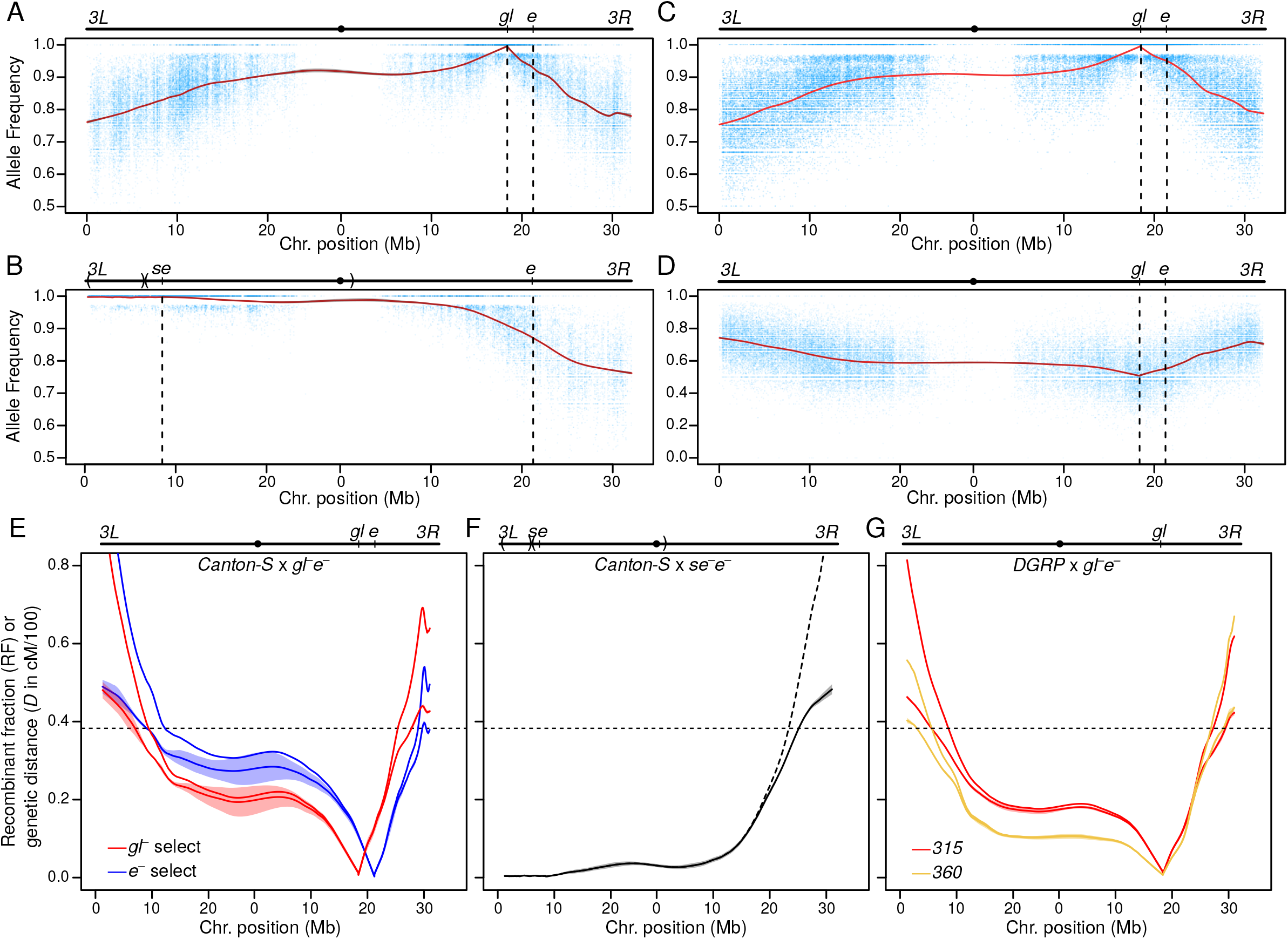
Recombinant fraction and genetic distance inferred from recombinant backcrosses with Chr. 3 markers. The per site (blue) and predicted (red line) allele frequency across the chromosome is displayed for four pools: A. Canton-S x *gl^−^ e^−^* with *gl^−^* selection (Table 1, row 6), B. Canton-S x *se^−^ e^−^* with se^−^ selection (Table 1, row 1), C. DGRP-315 x *gl^−^ e^−^* with *gl*^−^ selection (Table 1, row 12), and D. DGRP-315 x *gl^−^ e^−^* with *gl*^+^ selection (Table 1, row 13). For all marker selected pools, see Supplementary figure 3. Note that D has a different Y-axis scale. The positions of the double markers are marked in the chromosome schematics above and with vertical dotted lines in the plots. The gray interval (barely visible) around the predicted allele frequency mark the 95% confidence interval determined by the LOESS fit. For the Canton-S x *se^−^ e^−^* cross, large inversions are marked in chromosome schematic above with parentheses. E-G. Averaged recombinant fraction (solid curve) and genetic distance (dotted curve) from the selected loci. Interval around the recombinant fraction estimates represent the standard error across the replicates. Horizontal dotted lines represent recombinant fraction equivalent of 50cM after Kosambi transformation. E. Canton-S x *gl^−^ e^−^* crosses when *gl^−^* (red) or *e^−^* (blue) are selected. Note here that *gl^−^* pools were the female pools and *e^−^* pools were the male pools. F. Canton-S x *se^−^ e^−^* crosses with *e^−^* selection. G. DGRP-315 (red) or DGRP-360 (yellow) x *gl^−^ e*^−^ crosses. For the DGRP-315 cross, values are averaged across *gl*^+^ and *gl^−^* selection pools, while only *gl^−^* pools were sequenced for DGRP-315.

### AF decay (or the lack thereof) captures inversion induced crossover suppression

Interestingly, unlike the crosses with the *gl^−^ e^−^* double marker strain, the Canton-S x *se*^−^ *e*^−^ crossed produced AF curves that do not have a clear peak flanked by distal decay (Figure 3B). The AF on Chr. 3L where the selected locus *se* resides shows minimal decay with elevated level of AF extending into 3R, where it begins showing clear attenuation. Based on *e^−^* selected pool in the Canton-S x *gl^−^ e^−^* cross, the recombinant fraction between *e* and *se* is 0.368 equating to 47.0 cM (Figure 3E, blue line), but the estimates from both fly scoring and AF decay using the *se^−^ e^−^* marker strain indicate a substantially lower recombinant fraction of 0.238 equating to 25.8 cM (Table 1, Figure 3F). While the drastically shorter genetic distance of 3L may be due to bona-fide reduction in recombination rates along 3L in *se^−^ e^−^* strain, we suspect that this may be the result of a large inversion on this chromosome arm leading to the suppression of crossovers. We used Lumpy to infer the presence of structural variants for the whole genome sequencing; we indeed found at two large inversions on 3L: one from 0.25 Mb to 7.22 Mb and another pericentric inversion from 7.81 Mb to 2.08 Mb of 3R (Figure 3B and F), confirming our suspicion. The minor reduction in AF likely reflects the rate of rare double (or even number) crossover events which resolve the deleterious rearrangements caused by single (or odd number) crossover events within the inversion.

### Accounting for offsite viability effects that modulate AF decay

While the AF decay pattern is a product of crossovers between the selected and distal loci, it is also sensitive to loci with alleles that affect viability, body size, and fitness (which we will collectively refer to as viability effects), as such alleles can cause non-Mendelian contributions of individuals and DNA in the final pool. For example, in using double mutant marker lines like *gl^−^ e^−^*, the unselected locus may potentially have deleterious impact and affect the AF due to reduced number of homozygous mutant individuals. To evaluate how such loci will affect AF estimates, we first expand on the mathematical relationship between AF and recombinant fraction (*D*) by a secondary locus at site *l* that also modulates AF in addition to the selected locus at site *l*. For simplicity, we are removing the paternal contribution. The two loci have respective viability effect of *s*_*o*_ and *s*_*l*_ that individually (absent of other contributing loci) cause AF 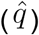 of:

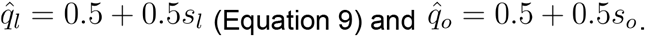

Positive and negative values of *s* mean that an allele contributes more and less in the final pool compared to the homologous allele, respectively, thus resulting in AFs that are higher and lower than the Mendelian expectation, respectively. Given that the distance between *l* and *o* is *D*_*lo*_, the AF at *o* is then:

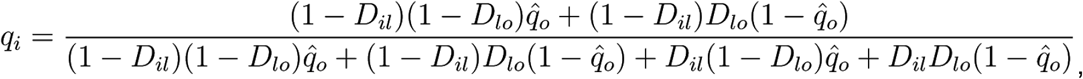

With the selection process at *l*, *s_l_* should always be 1 and 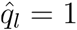, so the equation reduces to:

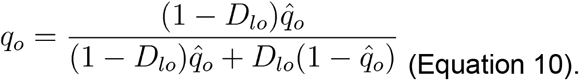

From this, it can be inferred that:

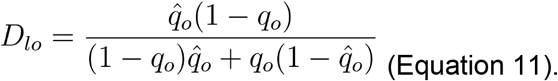

The AF at *i*(*q*_*i*_) will not only depend on its genetic distance from *l* (*D*_*il*_) and *o* (*D*_*i0*_) but also where it is positioned with respect to the two loci as its relative positions affect the recombinant allele combinations (Figure 4). First we will consider no crossover interference. When *l* is between *i* and *o*, i.e. **H**(*D*_*i0*_) = **H**(*D*_*il*_) + **H**(*D*_*l0*_)(Figure 4A),

**Figure 4.**
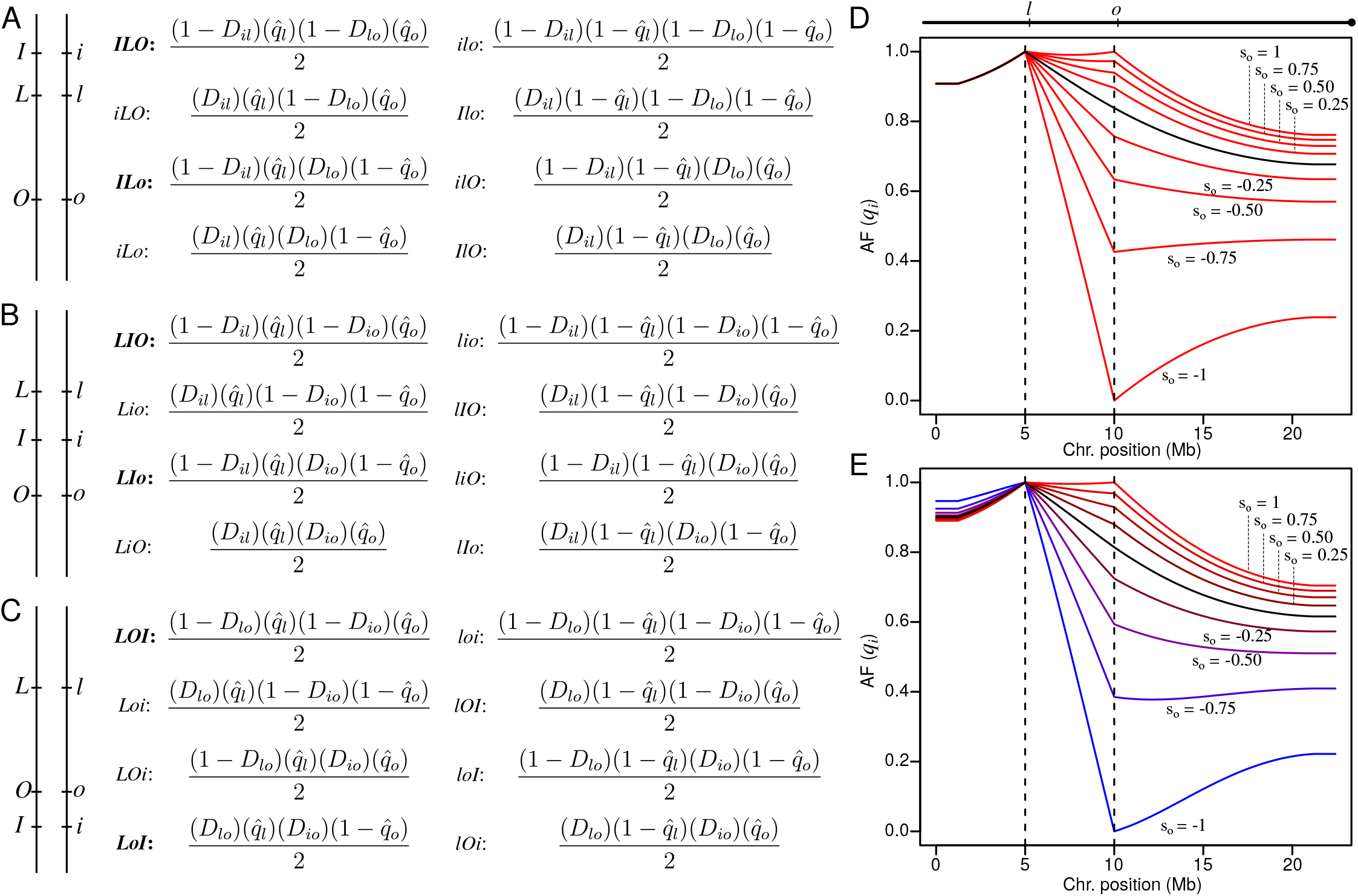
Effects of a secondary locus on allele frequency decay. In the marker selected pool, the selected locus (l), the offsite locus that affects viability (o), and any position along the chromosome (i), effectively create a three point cross. Upper- and lowercase letters of these loci differentiate the homologous alleles. A-C. Depending on the relative position of the three loci along the chromosome (schematics on the left), different allele combinations are produced at different frequencies (right). The frequencies of all possible allelic combinations are described by the equations which take into account of the fitness of the selected and offsite loci (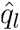 and 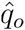, respectively). When 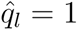, only the allelic combinations on the left column are incorporated in the marker selected pool and allelic combinations on the right will all equal 0. The combinations in bold are those that contain the uppercase I allele. Equations 10-13 are derived from dividing the sum of the allelic combinations in bold by the sum of the allelic combinations on the left for each three point cross. The frequencies of the different allelic combinations assume no interference. D. Allele frequency modulation given different values of *s*_*o*_ based on the X chromosome recombination rate, Haldane’s transformation, selected locus at 5Mb, and secondary locus at 10Mb. Black curve represents *s*_*o*_ = 0 (no secondary offsite viability effect). E. Same as D, but with Kosambi’s transformation and taking into account of interference with *C* = 2*D*. Colors are added to differentiate between the curves with different values of *s*_*o*_. Formula for how interference is incorporated into the allele frequencies can be found in supplementary figure 8.

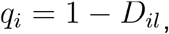

This simplifies to:

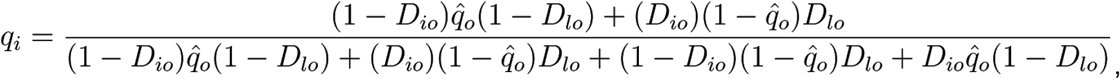

which indicates that as long as *s*_*l*_ = 1, *s*_*o*_ will have no effect on the allele frequency on the distal side of *l*. When *i* is between *l* and *o*, i.e. **H**(*D*_*l0*_) = **H**(*D*_*il*_) + **H**(*D*_*i0*_)(Figure 4B),

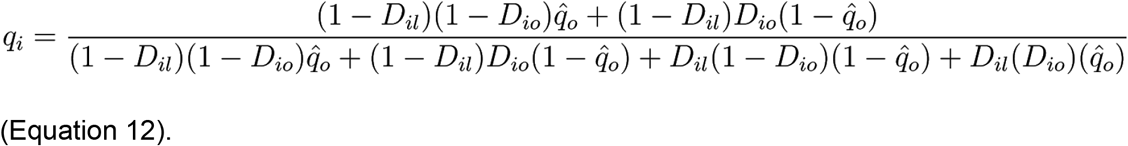

When is *o* between *l* and *i*, i.e. **H**(*D*_*il*_) = **H**(*D*_*lo*_) + **H**(*D*_*i0*_)(Figure 4C),

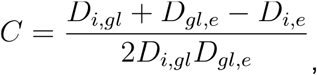

and reduces to:

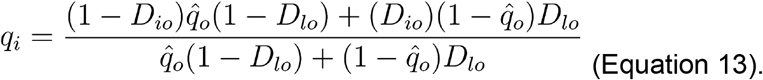

To account for interference, the double crossover components are multiplied with the coefficient of coincidence (2D for the Kosambi function) with single crossovers reciprocally increased (see Supplementary figure 8). To illustrate these relationships, we, again, used the recombination rate from the Recombination Rate Calculator(Fiston-Lavier *et al.* 2010) and varied the degree of *s*_*o*_. In both the presence and absence of interference, the AF is more sensitive to negative fitness impact at the secondary loci (*s*_*o*_ < 0). At extreme values of *s*_*o*_ (Figure 4D), local maximum and minimums can be observed, but at intermediate levels, the AF curves do not show obvious “kinks”. Unlike the curves in the absence of interference, *s*_*o*_ can modulate the the AF across the selected locus (Figure 4E). Unexpectedly, at higher *s*_*o*_, the AF is lower on the other side of *l*. The reason for this is that a positive *s*_*o*_ ensures more individuals lacking crossovers between *l* and *o*, which, in the presence of interference, results in increased probability of crossovers on the other side of *l* and therefore faster AF decline.

Given these relationships, it is then possible to solve for *D*_*il*_, *D*_*lo*_, and *D*_*io*_ when *s*_*o*_ and *q*_*i*_ are known. While *D*_*lo*_ can be deduced based on the Equation 11, we solve for *D*_*il*_ computationally using a root-finding algorithm since *D*_*il*_ cannot be easily isolated in the equations (see Material and Methods). While in our simulations *s*_*o*_ is known, in practice it needs to be determined empirically. This can be done by pooling and sequencing all BC1 individuals (without marker selection) and identifying regions with non-Mendelian rates. To emulate this, for the DGRP315 x *gl^−^ e^−^* crosses where we sequenced both positive and negative marker selected pools, we summed the AF at each site between the two, reasoning that the AF at the selected locus *gl* should then be Mendelian, and additional loci with viability effects (e.g. *e^−^*) will deviate from the Mendelian rates. Using this method, we indeed find a minor allele frequency deviation at *e^−^*, which is equivalent to (Supplementary Figure 9A). Correcting for this resulted in negligible changes in the recombinant fraction (Supplementary Figure 9B).

### Natural variation in recombination rate

Since recombination rate is genetic distance per physical distance, for regions to the left and right of the selected locus, the recombination rate is the negative and positive slope of the genetic distance curve, respectively. We converted the genetic distance to recombination rate for all the *gl^−^ e^−^* crosses by taking the slope of *d* in 500bp windows. We then compared between the recombination rate across the three lines, Canton-S, DGRP-315, and DGRP-360 (Figure 5A). As expected, the recombination rate is minimal across the pericentromeric region, but gradually increases away from the centromere. Overall, recombination rate increases toward the chromosome ends, followed by a drop off near telomeric ends, a pattern consistent with previous findings (Fiston-Lavier *et al.* 2010). Furthermore, our estimates are significantly correlated with past genome-wide estimates based on crossover breakpoint distribution by Comeron et al., 2012 (Comeron *et al.* 2012) (Pearson’s r = 0.4926, p-value < 2.2e-16) (Figure 5B). Reassuringly, the range of the rates are also similar, indicating that our method is not substantially over- or underestimating. Curiously, our estimates are on average significantly higher by 1.31-fold (paired Wilcoxon rank sum test p = 2.34e-06) (Figure 5C and Supplementary figure 10), and therefore produces a longer genetic map. By summing the genetic distances of windows within <50cM or the selected locus (red points in Figure 5B), estimates from our pools produced a genetic map of 99.18cM from 5.15 Mb on 3L to 29.2 Mp on 3R; in contrast, the same region is only 82.04 cM based on Comeron et al. To determine whether this inflation reflects overestimations resulting from the exponential growth of the mapping function at higher genetic distances, we limited the estimates to regions less than 15 cM away from the selected locus (Figure 5B and C). We similarly found significantly elevated rates (paired Wilcoxon rank sum test p = 1.062e-05), indicating that the transformation is not inflating the genetic distances and recombination rates (Figure 5B). Thus, the difference reflects bona-fide higher recombination rate likely due to environmental differences of the crosses. For example, we raised and crossed our flies at 25°C instead of the 23°C by Comeron et al, 2012 and higher temperature is known to increase recombination rate (Stern 1926).

**Figure 5.**
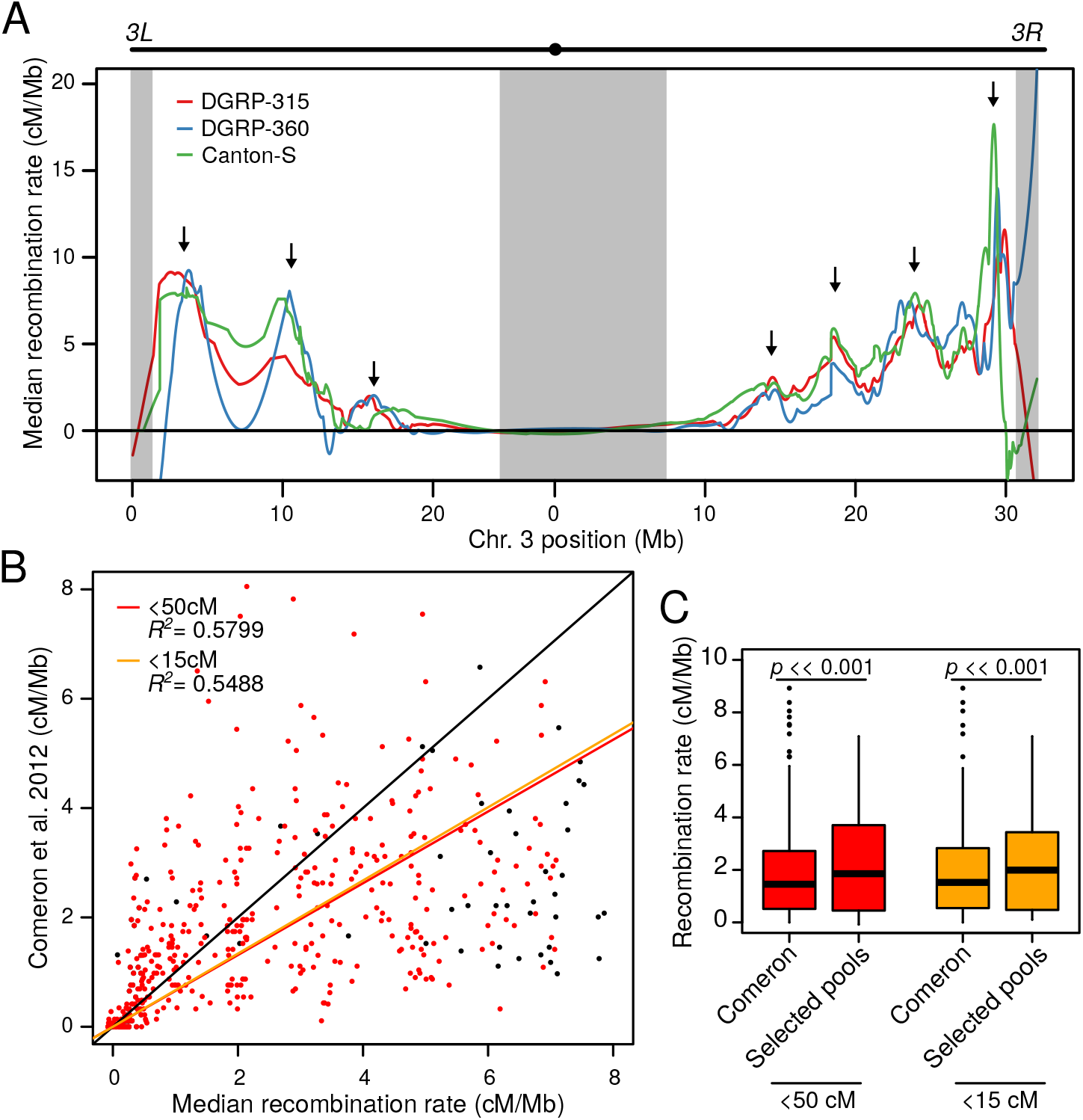
Recombination rate estimates in wild-derived strains. A. The slope of the genetic distances in Figure 3E and G are taken in 500bp windows to estimate the Chr. 3 recombination rate for Canton-S, DGRP-315, DGRP-360. The averages across all replicates plotted. Arrows point to regions of the genome with elevated recombination rate in all three lines. Gray boxes mark regions of the genome with high repeat content and low SNP density. B. Correlation of recombination rates in A averaged with previous genome-wide estimates by Comeron et al 2012. Each point is the estimates of a 100 kb window. Red points are windows that are less than 50 cM away from the selected locus based on genetic distance in Figure 3E and G. Black diagonal marks the identity line, red line marks the regression using only the red points, and yellow line marks the regression using points that are less than 15 cM away from the selected locus. C. Boxplots depict the distribution of the estimated recombination rates from the two methods, depending on whether the 50 cM or 15 cM cutoffs were used.

Interestingly, we see multiple spikes of recombination rate across all lines with three clear peaks on 3L and four on 3R that are found across all three lines (Figure 5A). While these could reflect loci with viability effects of the *gl^−^ e^−^* chromosome, we find this unlikely given the absence of strong offsite fitness effects across this chromosome (Supplementary figure 9). Therefore, these shared peaks and valleys likely reflect either regions of the genome with robust rates that are insensitive to genetic background, or regions with modulating recombination rate due to a common environment. Despite having similar overall patterns, the strains also show notable differences. On 3L, the recombination rate of DGRP-360 drops to 0 at ~8Mb, while the other two strains show less substantial dips. On 3R DGRP-360, consistently show lower rates to up to 22Mb, which resulted in a shorter genetic distance (Figure 3G).

### Inferring crossover interference from reciprocal selection of double markers

Typically crossover interference, usually expressed 1 − *C* where *C* is the coefficient of coincidence, is estimated from three point crosses and the observed number of double recombinants is compared to the expected number of double recombinants. Given Equation 2’, it can also be estimated when the recombinant fractions between the three loci are known. In our crosses using Canton-S, we sequenced pools of both *gl^−^* and *e^−^* selection, generating chromosome-wide recombinant fractions with respect to these two loci. With these two curves, we can estimate the coefficient of coincidence, as for any site *i*, we have the genetic fractions between *gl* and *i*(*D*_*i,gl*_), e and *i*(*D*_*i,e*_), and *gl* and *e* (*D*_*gl,e*_). Therefore to the left of *gl*, between the two, and right of e, we can solve for *C* with the following, respectively:

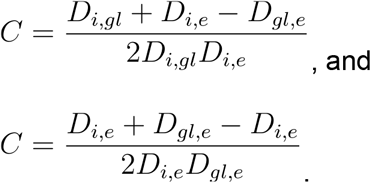

Because of the denominator, regions close to loci with small will yield nonsensical results. We therefore, only estimate the interference beyond ±2.5Mb from the markers, which also precludes estimating between *gl* and *e*, as they are only 2.8 Mb apart. To the right of *e*, C increases distally towards the 3R telomere. To the left of *gl*, while C also increases distally, toward 3L telomere, a sharp peak ~5Mb away from the selected locus is also observed. Unfortunately, these Canton-S samples were the most variable of the crosses, therefore it is difficult to determine whether this, along with other peaks and troughs, are bona fide regions of low and high interference, respectively. Nevertheless, in comparison to the coefficient of coincidence with the Kosambi function, we see that *C* remains low for longer but recovers towards 1 more precipitously, suggesting that interference remain strong near crossovers for a longer stretch, but weakens quickly distally.

**Figure 6.**
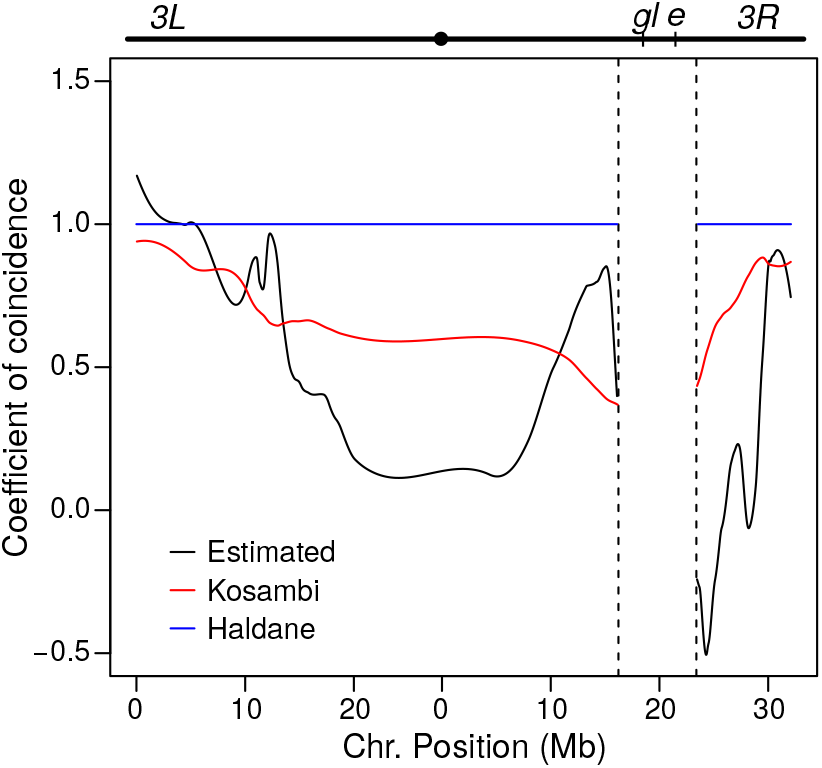
Estimating coefficient of coincidence from reciprocal marker selection pools with double markers. In the Canton-S x *gl*^−^ *e*^−^ crosses, males were selected by *e*^−^ and females were selected by *gl*^−^. The chromosome-wide recombinant fractions from these two selected pools can be used to estimate the coefficient of coincidence across the chromosomes with respect to these two loci (black line). Estimates near the selected locus (within 5 Mb) as the values approach their limits. For reference the coefficient of coincidence based on the Haldane and Kosambi mapping functions are in blue and red respectively.

## DISCUSSION

### Broad range recombination rate estimation using allele frequency in marker selected pools

We demonstrated theoretically and empirically that allele frequency decay around a selected marker in a recombinant backcross can be used to infer near chromosome-wide recombination rate. Instead of genotyping individuals for recombinant breakpoints, this approach relies on pooled sequencing of large numbers of marker-selected individuals from recombinant backcrosses. Since the allele frequency surrounding the selected locus which decays proportionally with the recombinant fraction, the genetic distance can be determined using mapping functions, and the slope will then approximate the recombination rate. Since each pool can provide broad range recombination rate estimates, this method substantially reduces the number of library preparations needed for chromosome-wide estimates. Previously, hundreds to thousands of individuals needed to be sequenced or genotyped to capture a comprehensive spread of recombinant breakpoints from which chromosome-wide recombination rates are inferred, here we used as little as two libraries for near chromosome-wide estimates. Using this approach we inferred the recombination rate of Chr. 3 of three wild-derived lines, and demonstrated how it can be used to infer crossover interference.

As mentioned in the introduction, Singh et al. 2013, similarly used allele frequency to estimate crossover frequency; recombinants between garnet and scalloped that carry either one of the two markers are pooled to estimate the fine scale recombination rate between the two (Singh *et al.* 2013). The method presented here is fundamentally different from Singh et al. 2013, in that recombinants are not explicitly targeted for selection, and as such the allele frequency decay is directly reflecting the recombinant fraction (Equations 4 and 5). When only recombinants between two loci are selected, the allele frequency decay is effectively restricted to estimates of recombinant fraction between the two loci. As demonstrated by Gilliland 2015, the approach used by Singh et al. 2013 to measure allele frequency and recombination rate produces estimates that are strongly anti-correlated with the heterogeneous window sizes (Gilliland 2015). With our method, we purposefully avoid estimating recombinant fraction at small window sizes. As demonstrated by our simulations and pointed out by Gilliland 2015, the variability of read counts at individual sites are too large for fine scaled estimates.

### Methodological limitations and workarounds

Given that our method does not directly identify crossover breakpoints, the downside of the high-throughput and cost-effectiveness is lowered resolution. In addition there are several limitations to the method. First, a visible marker is obviously important for the selection process. However, given Equation 9, it is possible to estimate recombination rate around a locus with strong deleterious viability effects instead of a marked locus under selection, even if the effect is not fully penetrant (−1 < *s*_*l*_ < 1) (Supplementary figure 11). The closer to 0 *s*_*l*_ is, the sampling error around the read counts is expected to increase resulting in noisier estimates.

Second, the conversion from recombinant fraction to genetic distance with mapping functions requires assumption about crossover interference. The Haldane, Kosambi, and other mapping functions have explicit assumptions about the extent (or lack) of interference, which accordingly affect the genetic distance conversion, with higher crossover interference producing shorter genetic maps. Nevertheless, we presented an extension of our method to infer crossover interference; by reciprocally selecting for markers in a double marker strain, we were able to estimate the coefficient of coincidence across Chr. 3 with respect to the two markers. Alternatively, since the mapping functions have negligible effects for smaller values of recombinant fractions (< 10), multiple markers (if available) along the chromosome can be used to estimate recombination rate in small intervals. While this significantly increases the number of crosses needed, the number of libraries generated is still at least one order of magnitude less than sequencing individuals. As a corollary, mapping functions break down when the recombinant fraction approaches 0.5. Here we explicitly restrict the Kosambi mapping function to a genetic distance of 50 cM on either side of the selected loci, which translates to recombinant a total measurable fraction of 0.762 (0.381 on either side). Again, to extend the genetic map, multiple markers along the chromosome can be strategically chosen to encompass the entirety of the chromosome. Down the line, improved understanding of crossover interference in a genomic context will allow for more accurate conversion between the recombinant fraction to genetic distance.

Third, this approach is sensitive to the SNP density. Based on our simulation, this approach is robust even when the SNP density is lower than 1 site per 1000bp. While intraspecific strain differences are likely higher, polymorphisms are not evenly distributed across the chromosome. The SNP density drops rapidly around the pericentromeric and telomeric regions, which has resulted in increased error rates around those regions. While the decrease in SNP density is particularly problematic in window based AF estimates, our usage of the LOESS fit alleviates this issue on at least two fronts: the smoothing is conducted based on the number of sites instead of genomic windows and the reduced pericentromeric SNP density on 3R is “stabilized” by increased density across the centromere on 3L. We note, additionally, that reduced SNP density also poses a challenge when inferring breakpoints, since the precise location of haplotype changes will be difficult to pinpoint.

Lastly, as we demonstrated, offsite viability effects can modulate the allele frequency decay around the selected locus. Such viability effects can result from alleles that induces lethality or reduces body size, both of which will change the allele frequency in the DNA pool. The former is similarly problematic for genotyping and breakpoint inference in individuals, since it reduces the number of recoverable recombinants at specific loci, but are typically ignored. We analytically showed how additive effects of offsite viability effects can be accounted for, provided that the extent of the fitness reduction (*s*_*o*_) can be determined. To estimate *s*_*o*_ without additional experiments, we simply summed the allele frequency of both the positive and negative marker selection pools that originated from the same cross which effectively removes the peak and the effect of the selected locus; the remaining elevations and drops that deviate from Mendelian ratios are then regions with viability effects.

### Applications beyond *Drosophila*

While this study focused on *D. melanogaster*, this method is readily applicable in other organisms, particularly for model species with a wealth of phenotypic markers readily available like *C. elegans* (Greenwald 2016). As discussed above, for species with longer genetic maps, multiple markers will need to be used to capture the entirety of the chromosome, thus increasing the number of crosses and libraries. Since each marker selected pool only reflects recombination rate on one chromosome, species with higher chromosome numbers will require more libraries and crosses overall. Notably, markers from different chromosomes can be selected at the same time in any given cross allowing for estimates on different chromosomes in parallel. However, this may also introduces potential epistatic effects between markers which may potentially modulate allele frequencies of the chromosomes. As with typical recombinant crosses, the sensitivity of the method depends on the number of BC1 individuals (Supplementary figure 1). While we recommend over one thousand individuals to be collected, for organisms where large number of offsprings are unfeasible, the estimates from marker selected pools will be similarly underpowered as counting the number of recombinant individuals between markers, but with the added benefit of requiring only one marker. For species like mice where pulverizing individuals is impractical, care must be taken when pooling tissues to minimize variation in tissue size and the resulting DNA contribution.

Overall, sequencing marker selected pool is a simple, cost-efficient, and widely applicable approach to estimate near chromosome-wide recombination rates that require orders of magnitude fewer libraries than previous approaches.

## MATERIALS AND METHODS

### MarSuPial

The analytical and statistical methods described below can be found in the MarSuPial package found in KW’s github page (https://github.com/weikevinhc). It is an R package with tools to analyze and simulate read count data from marker selected pools.

### Conversion and simulation of recombinant fraction from recombination rate function

To convert recombination rate (from Recombination Rate Calculator (Fiston-Lavier *et al.* 2010)) to genetic distance centered at the selected locus (e.g. *w*) to, we integrated the quadratic formula from the selected locus to every position on the chromosome. For example, the genetic distance between w (at position X:2,684,632) and position in megabses is:

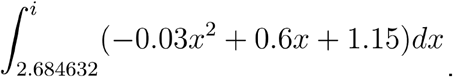

To convert genetic distance to recombinant fraction using the Haldane and Kosambi transformations, we used the functions 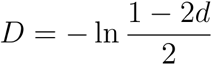 (Haldane 1919) and 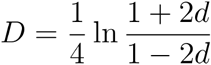 (Kosambi 1943), which are derived from Equations 1 and 3, respectively. Since the formula from the Recombination Rate Calculator are based on r5 of the *D. melanogaster* genome, we used the r5 instead of r6 coordinates for the genes in the simulations. The remaining analyses were all based on the r6 reference. For read count and allele frequency simulations, based on the chromosome wide recombinant fractions, the allele frequency with paternal contribution (*q*) is determined at 1000bp intervals. At each of these positions, a binomial draw is conducted with a frequency *q* and sample size of 2*n* where *n* is the number of individuals in the pool and a Poisson draw is conducted with the desired read depth as the sample size. These two draws simulate the sampling involved in allele counts in the pool and read counts from sequencing, respectively. A second binomial draw is then conducted with the binomial-drawn frequency and Poisson-drawn read depth, to simulate the allele-specific read counts at the site. To determine the effects of varying read depths and pool (sample) sizes, we generated 20000 chromosome-wide trials for each condition; summary statistics are then determined from the trials.

### Predicting and smoothing the allele frequency with linear regression and LOESS

Once AF at differentiating sites are determined from read counts (simulated or real) across the chromosome, sites within 500kb windows are then used for a weighted linear regression with the R function lm() where the weight of each site is the coverage of that site. The AF of the window is then determined for the midpoint of the window using the slope and y-intercept of the linear regression. Given the linear regression, AF can also be predicted for any point within the window. AF in 500kb windows are determined every 100kb, resulting in overlapping sliding windows.

For LOESS fit, we use the R function loess(), to fit two curves on either side of the selected locus, which prevents smoothing of the expected peak/trough. The AF at each site is weighted by the coverage. To “anchor” the LOESS fit, we include an additional point at exactly the selected locus with the expected AF (0, 0.5, or 1, depending on the selection and/or chromosome), with a weight of 1000000, ensuring that the fit reaches or begins with the expected AF. Given that the two LOESS fit to the left and right of the selected locus are stored as R objects, the predict() function on these objects allows for estimates of the AF and standard error for any position across the chromosome.

### Genetic distance and recombination rate from allele frequency

Given the LOESS-predicted allele frequency chromosome-wide in 500bp windows for each cross, we removed the paternal contribution and inferred the recombinant fraction using Equation 6. The only exception is the DGRP315 x *gl^−^ e^−^* pool where the *gl*^+^ individuals were selected to which Equation 7 was used to obtain the recombinant fraction. We then applied the mapping functions (Equations 1 and 3) to get the genetic distance at each 500bp window. Recombination rate (*r*) at position *i* bp was then derived taking the positive or negative slope between the windows:

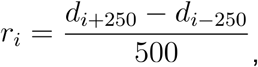

depending on whether *i* was to the left or right of the selected locus, respectively. Summary statistics for genetic distances and recombination rates (median, standard deviation, etc) were estimated from replicate and/or different sexed pools from the same cross. For the the Canton-S x *gl^−^ e^−^* crosses, the reciprocal marker selection produced two different sets of genetic distances from either glass or ebony. However, once converted into recombination rate, the reciprocals were then treated as replicates, since recombination rate is expected to be unaffected of the marker selected.

To convert the publically available recombination rate (in 100kb windows) by Comeron et al 2012 to genetic distance, we multiplied the recombination rate in each window by 100kb.

### Fly stocks, maintenance, and collection

The double marker lines were ordered from Bloomington Drosophila Stock Center, stock numbers BL1669 (*e^−^ se^−^*) and BL507 (*gl^−^ e^−^*). DGRP-315 and DGRP-360 are gifts from Dr. Grace Yuh Chwen Lee. All stocks and crosses were raised on standard molasses food at 25°C incubator. For the crosses, 4-8 day old virgin females were mated with the marker strain males and F1 virgins were then collected. For each recombinant cross, over 40 F1 virgins were collected and backcrossed to the marker males in vials of 5-8 virgin females to 8-10 males. To avoid overcrowding, vials were transferred every 2-3 days and after adults began to emerge, flies were cleared and scored daily. Every vial is collected for ten days to ensure that genotypes that may introduce developmental delays will not be underrepresented. Sexed and genotyped flies were maintained in fresh vials for 3-5 days prior to freezing.

### DNA extraction and library preparation

Frozen flies of the desired genotypes were pooled, pulverized with sterilized mortar pestle in liquid nitrogen, and then transferred to 50 mL falcon tubes. After adding 15mL of Cell Lysis Solution from Qiagen (Catalogue No.158908), samples were incubated at 65°C for 4hrs and vigorously shaken every hour. 75uL of ProteinaseK (Catalogue No. 19131) was then added and incubated at 55°C overnight. 600uL of the sample were then passed through the columns from the DNeasy kit (Catalogue No. 69506). The columns were then processed in accordance with the kit protocol. DNA for parental lines were extracted from 5 females using the same kit. The resulting DNA were fragmented to 550bp using the Covaris sonicator and libraries were made with the Illumina Truseq DNA Nano kit. Library quality were determined with the Bioanalyzer at the Functional Genomics Laboratory at UC Berkeley and samples were sequenced using the Illumina HiSeq 4000 machine at the Vincent J. Coates Genomics Sequencing Laboratory at UC Berkeley. Coverage of each cross can be found in Supplementary table 1.

### DNA-seq processing, genotyping, and filtering

Demultiplexed paired-end reads were mapped to the *D. melanogaster* genome r6.12 (Thurmond *et al.* 2019), using bwa mem (v0.7.15) on default settings (Li and Durbin 2009). Raw reads for the DGRP strains were downloaded from SRA. We removed duplicates using Picard tools (v2.18.14) (“Picard Tools - By Broad Institute”), and merged the parental strains with the crosses using Samtools (v1.5) (Li *et al.* 2009) to allow them to be genotyped together with GATK HaplotypeCaller (v3.8) (McKenna *et al.* 2010). By default, HaplotypeCaller only outputs sites where at least one sample has a non-reference variant. This is particularly problematic if samples were genotyped individually, since many of the sites will be deemed as homozygous reference and unreported, particularly around the selected locus in our selected pools. By genotyping the crosses together with the parents, we are ensuring that all informative sites are reported, because at least one of the two parents will be homozygous for the non-reference allele at informative sites. To filter for informative sites, we used bcftools (Li 2011)(v1.6) to first isolated the parental strains and retained only SNP sites where the parental strains have different homozygous alleles with both genotype quality of > 30 (Supplementary Table 2). In the crosses, all sites other than the informative sites are removed. No genotyping filter is applied on the crosses since many of the sites in the crosses will have intermediate allele frequency that are difficult to genotype. However, sites overlapping repeats and sites with coverage <0.5x and >1.5x of the chromosome average are removed as to avoid copy number variants. The coverage and number of SNP sites after filtering can be found in Supplementary table 1. The allele-specific read counts can be determined from the AD field in the vcf files (Danecek *et al.* 2011).

### Inversion identification

For structural variant calls, we used the smoove wrapper for Lumpy (v0.2.13) after aligning *e se* to the reference genome (Layer *et al.* 2014). We identified large structurals variant over 1Mb within and between Chr 3L and 3R.

### Removing offsite viability effects with root finding algorithm

Given the complex equations for the offsite viability effects, we use root finding algorithms to solve for *D*_*il*_ instead of isolating it from the equation. Since 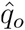 and *D*_*lo*_ can be predetermined, and *D*_*io*_ can be substituted with *D*_*il*_ in accordance with Equation 2, we are left with formulae with only *D*_*il*_ as the variable to solve. See supplementary figure X to see the formulae. We use the root finding function uniroot.all() from the package rootSolve in R to closely approximate the solution (Soetaert and Herman 2009). Note in some instances, more than one solution is possible, but usually only one is within reasonable range (between 0 and 0.5).

