## Supplementary material for "The theory and practice of measuring broad-range recombination rate from marker selected pools"

Supplementary table 1. Genotyping and sequencing summary for the recombinant backross pools

| Wild type strain (F) | Marker strain (M) | Sex | Selected phenotype | 3L | | | 3R | | |
|---|---|---|---|---|---|---|---|---|---|
| | | | | Before filtering* | After filtering | Mean Coverage | Before filtering* | After filtering | Mean Coverage |
| | | F | | 332877 | 12111 | 26.02 | 309530 | 10269 | 26.21 |
| Canton-S | *se', e'* | M | *se-* | 332877 | 12359 | 30.48 | 309530 | 10294 | 30.66 |
| | | F | | 332877 | 12363 | 32.57 | 309530 | 10451 | 33.01 |
| Canton-S | *se', e'* | M | *se-* | 332877 | 12553 | 30.10 | 309530 | 10429 | 30.41 |
| | | F | *gl-* | 332877 | 22664 | 27.54 | 309530 | 19936 | 27.79 |
| Canton-S | *e-, gl-* | M | *e-* | 332877 | 23244 | 26.86 | 309530 | 19661 | 26.78 |
| | | F | *gl-* | 332877 | 22954 | 29.34 | 309530 | 19974 | 29.49 |
| Canton-S | *e-, gl-* | M | *e-* | 332877 | 23722 | 32.96 | 309530 | 20419 | 33.09 |
| | | F | *gl-* | 460065 | 88201 | 17.82 | 434317 | 89253 | 17.60 |
| DGRP-360 | *e-, gl-* | M | *gl-* | 460065 | 86237 | 17.18 | 434317 | 86421 | 16.79 |
| | | | *gl-* | 460065 | 106649 | 20.12 | 434317 | 99149 | 20.00 |
| | | F | *gl+* | 460065 | 107223 | 20.92 | 434317 | 100570 | 20.56 |
| | | | gl- | 460065 | 107268 | 22.43 | 434317 | 101121 | 22.23 |
| DGRP-315† | *e-, gl-* | M | *gl+* | 460065 | 108374 | 21.80 | 434317 | 101887 | 21.69 |

*These numbers are the number of sites reported by GATK HaplotypeCaller. Samples genotyped together will have the same numbers

Supplementary table 2. Informative sites for recombinant backcrosses

| | 3L | | 3R | |
|---|---|---|---|---|
| | Before GQ filter | After GQ filter | Before GQ filter | After GQ filter |
| Canton-S x gl- e- | 264193 | 25927 | 247374 | 22219 |
| Canton-S x se- e- | 264193 | 13992 | 247374 | 11546 |
| DGRP-315 x gl- e- | 318869 | 121080 | 280881 | 111358 |
| DGRP-360 x gl- e- | 329834 | 100259 | 313045 | 100962 |

Supplementary table 3. Heterozygosity in parental lines

| | No. of Heterozygous Sites | |
|---|---|---|
| | 3L | 3R |
| se- e- | 5385 | 5854 |
| gl- e- | 5412 | 6232 |
| Canton-S | 78765 | 73798 |
| DGRP-315 | 13708 | 13616 |
| DGRP-360 | 38853 | 28922 |

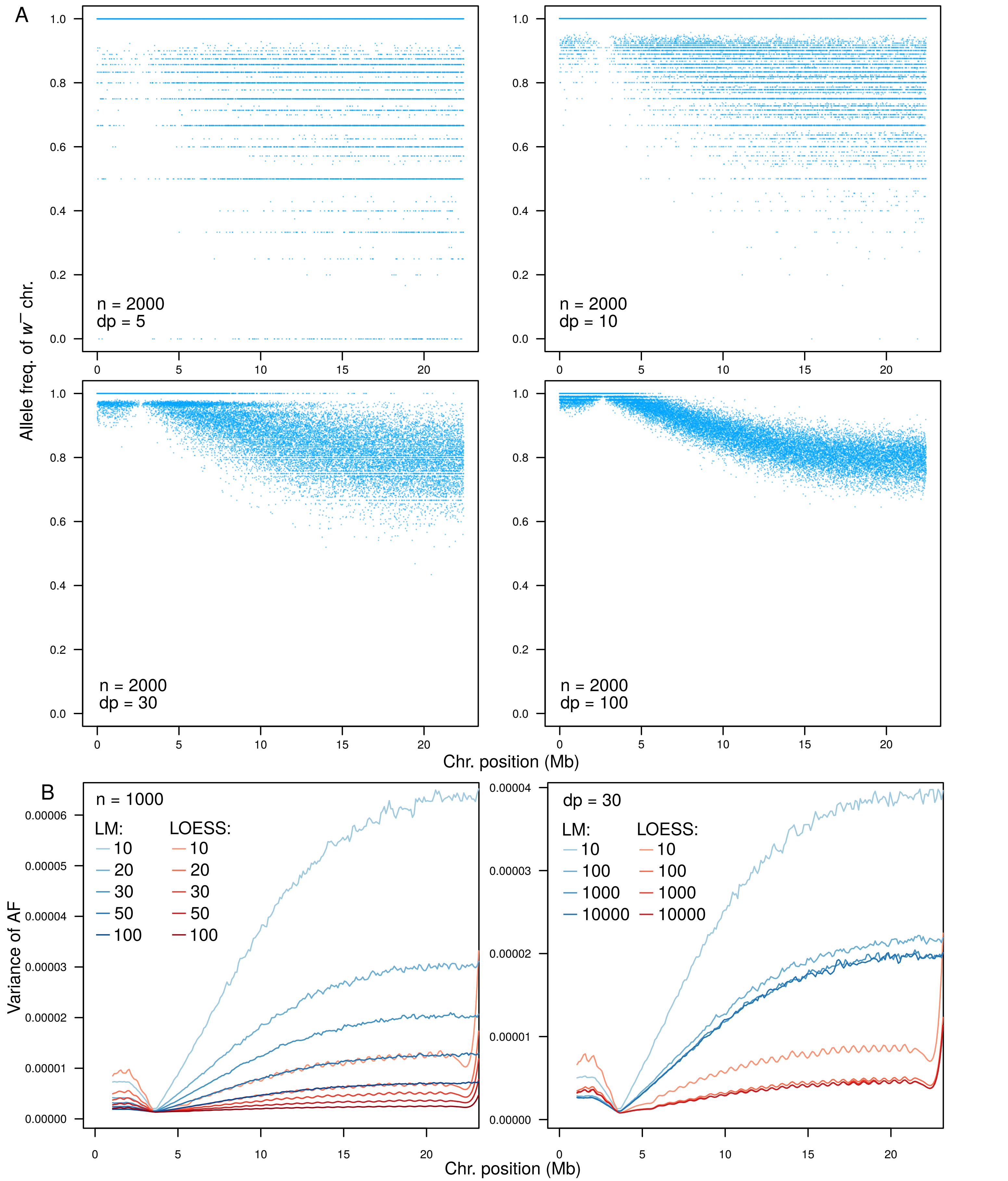


Supplementary Figure 1. Allele frequency and read count simulations based on $w^-$ selection with different sequence depths (dp) and pool sizes (n). A. Allele frequency inferred from read counts at individual SNP sites with different sequence depth. B. AF is estimated from linear regressions (LM) in overlapping sliding windows or local regressions (LOESS). Variance of the AF estimated from 10000-round Monte Carlo simulations is plotted given different sequencing depths (left), and pool sizes (right).

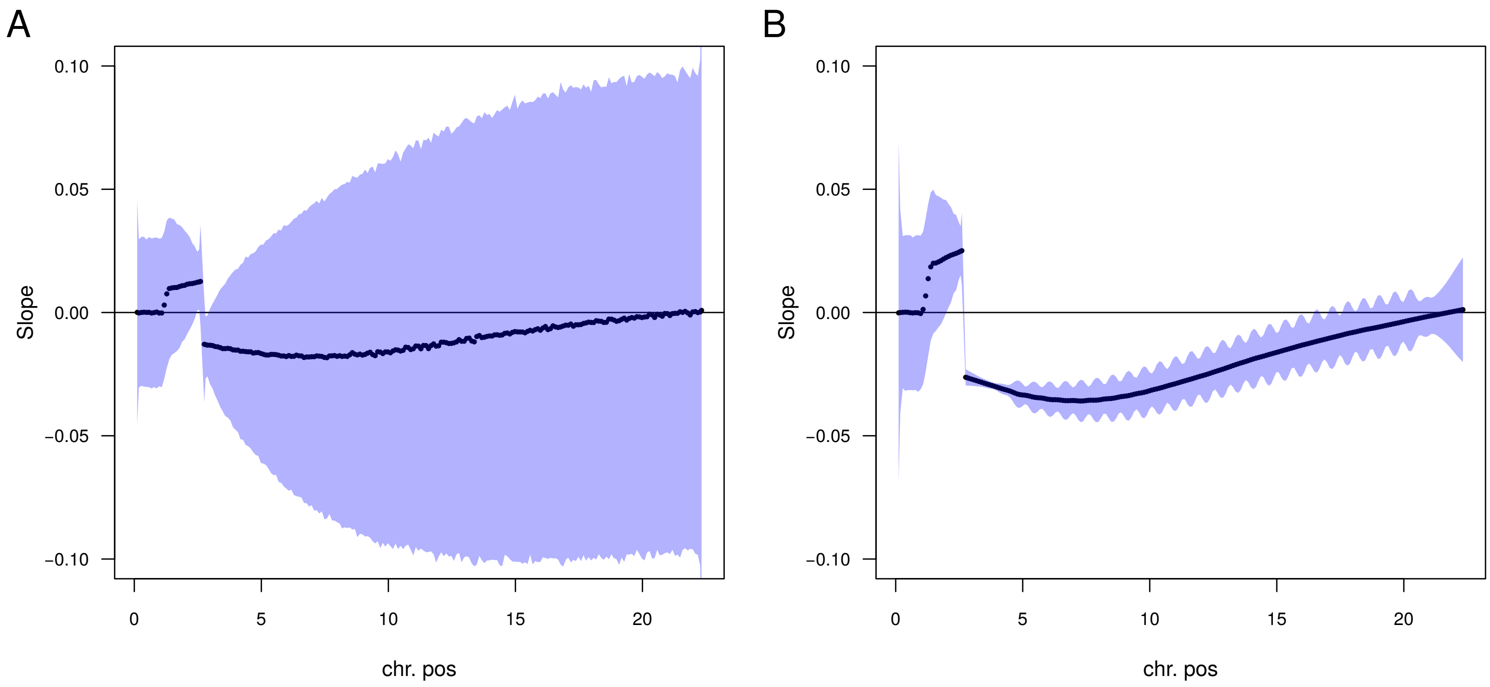


**Supplementary Figure 2**. Based on simulations with n = 1000 and dp = 30, the slopes from sliding window linear regressions (A) and LOESS fit (B) are plotted in black . The blue area represents the 95% confidence interval of the etimates.

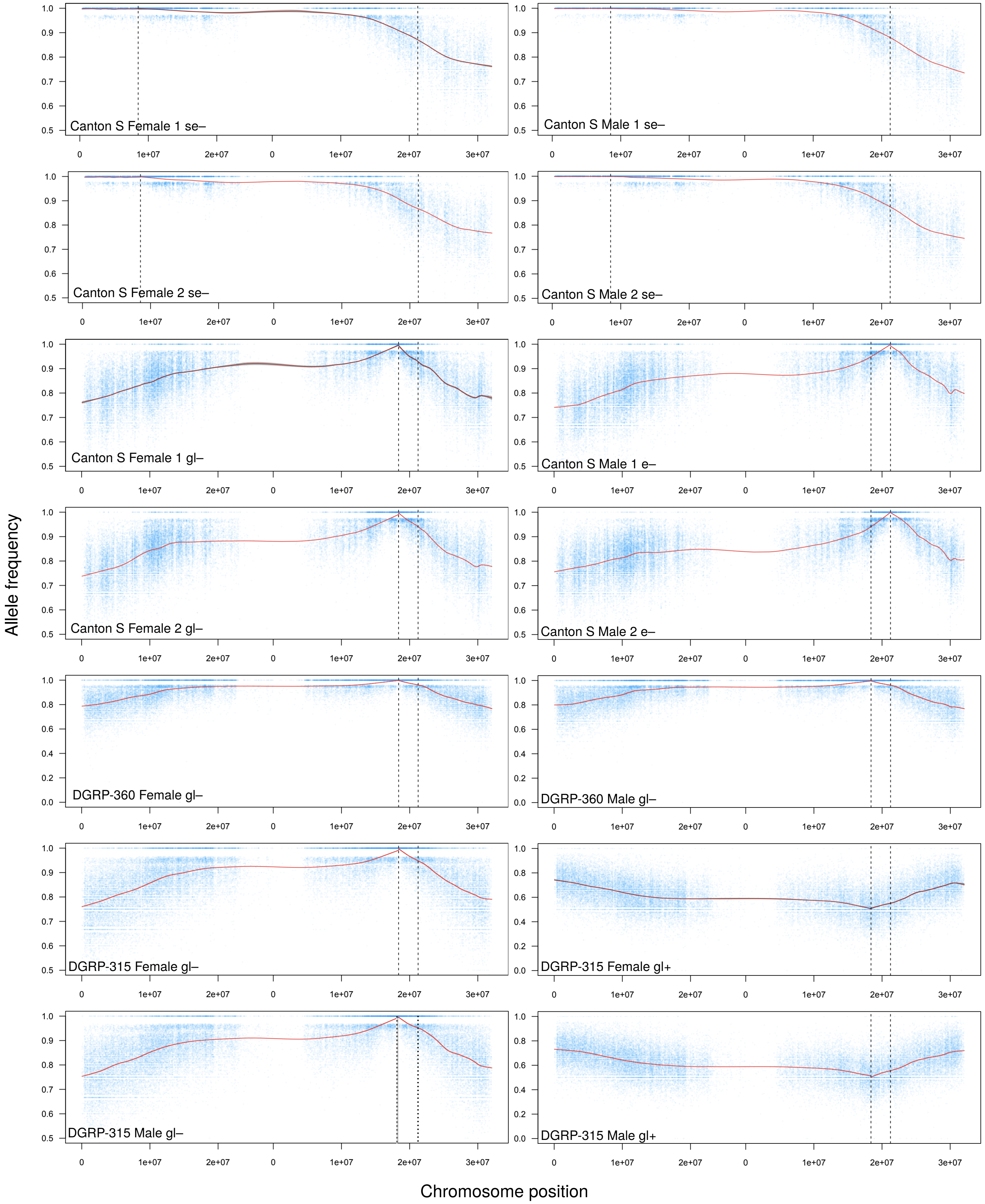


Supplementary Figure 3. Allele frequency estimates for all marker selected pools in Table 1. Blue dots represent allele frequency at individual informative sites. Red lines represent the LOESS estimated allele frequency.

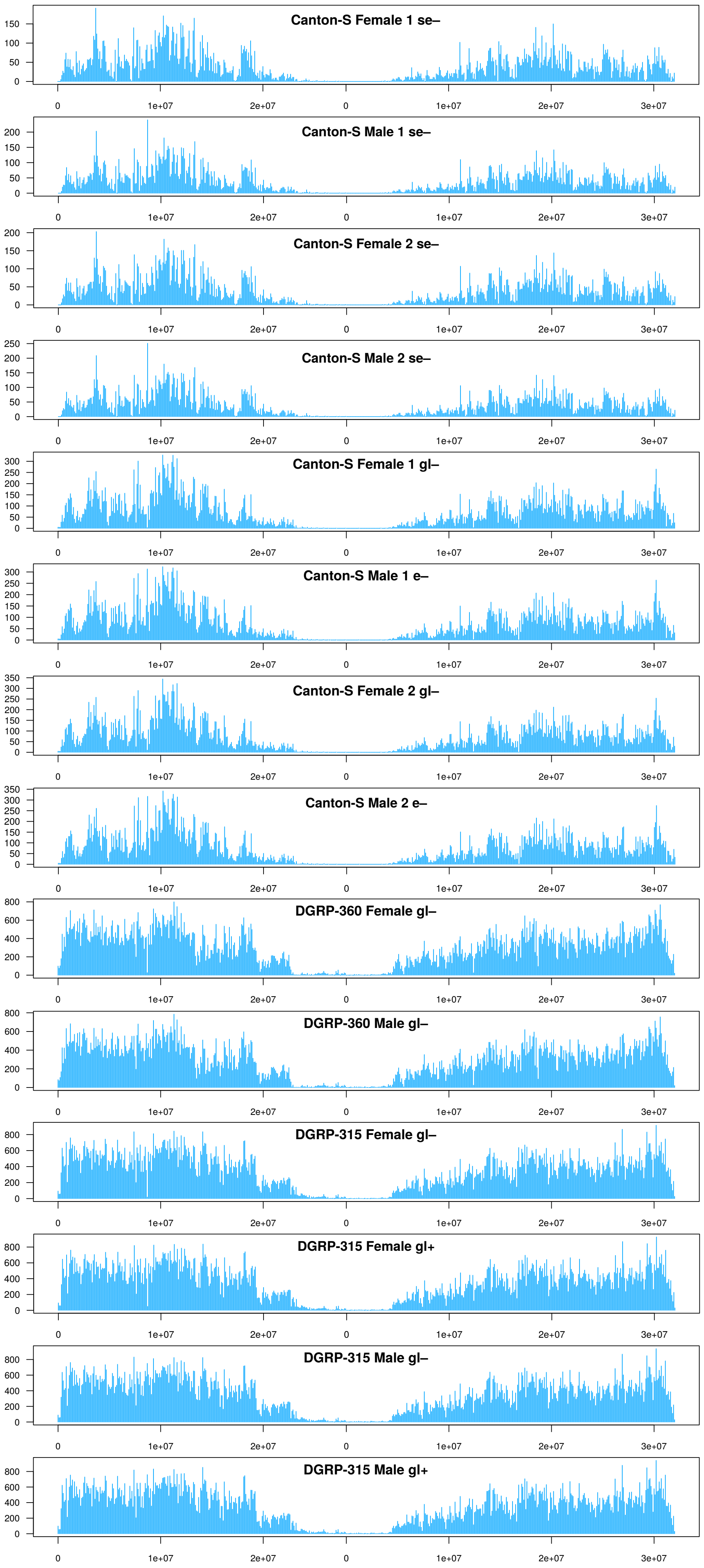


**Supplementary Figure 4.** Distribution of Informative SNP sites in backcrosses across Chr. 3L and 3R. Note that the Y-axes are not the same across plots. The plots are arranged in the same order as Table 1.

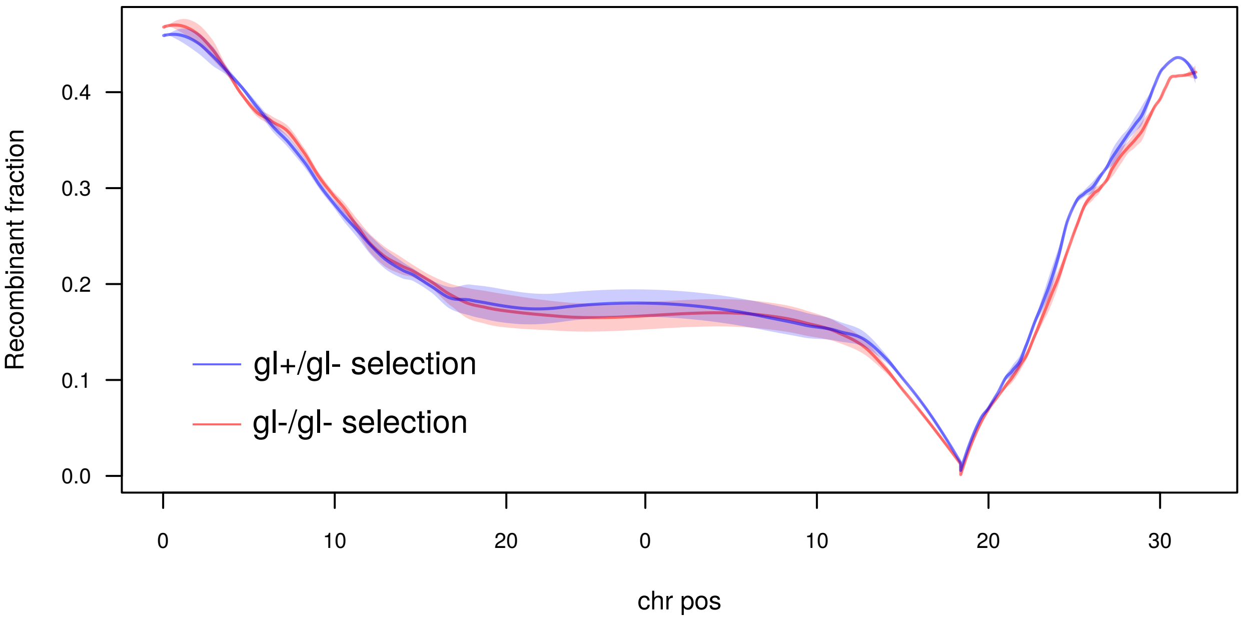


**Supplementary figure 5**. Positive and negative selection pools from the same DGRP-360 x gl- e- cross.

A

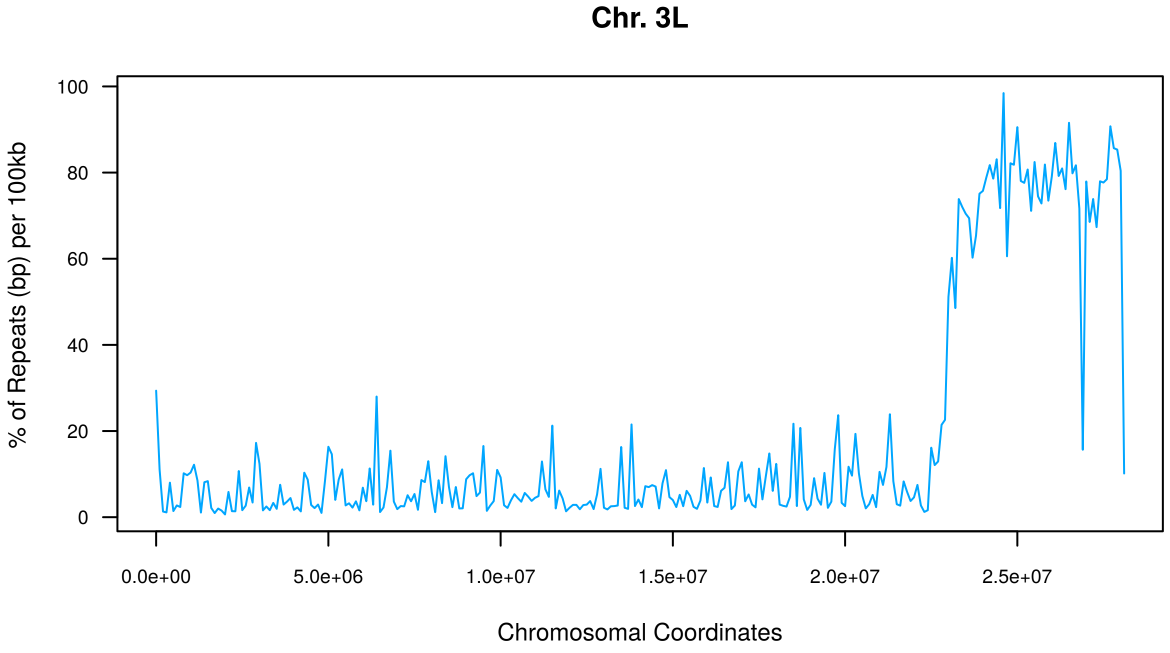


B

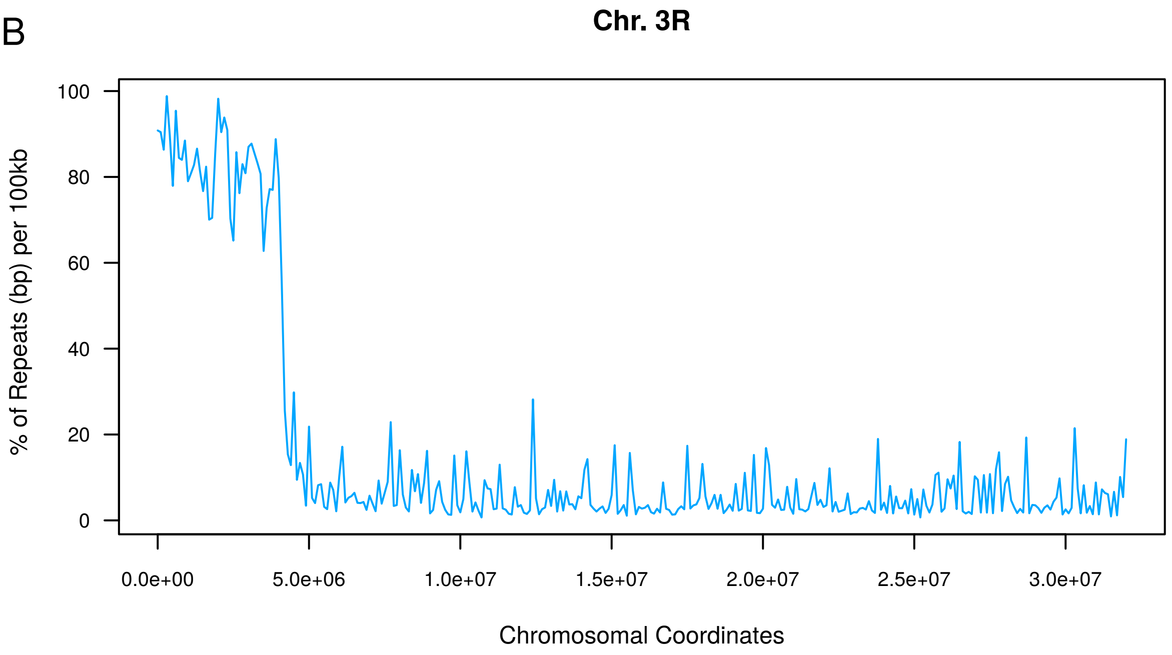


**Supplementary figure 6.** Repeat density on chromosome 3L (A) and 3R (B) of the reference genome (r6.12).

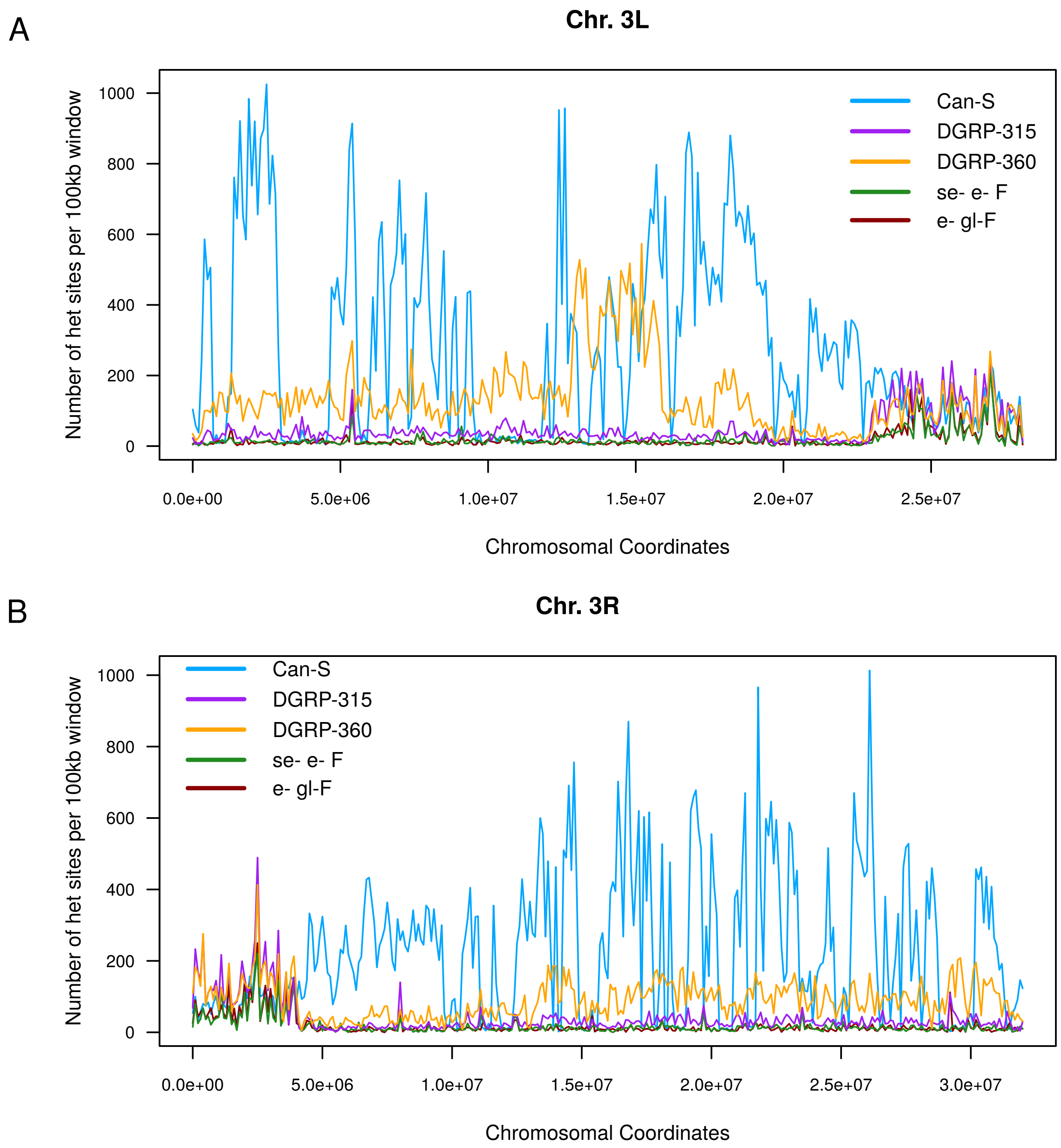


**Supplementary figure 7.** Heterozygosity in parental lines for chromsomes 3L (A) and 3R (B).

A

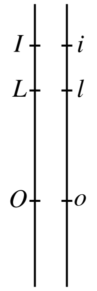


***ILO***: $\dfrac{(1-D_{il})(\hat{q}_l)(1-D_{lo})(\hat{q}_o)}{2}$

*ilo*: $\dfrac{(1-D_{il})(1-\hat{q}_l)(1-D_{lo})(1-\hat{q}_o)}{2}$

*iLO*: $\dfrac{(D_{il})(\hat{q}_l)(1-D_{lo})(\hat{q}_o)\left(\dfrac{1-D_{lo}C}{1-D_{lo}}\right)}{2}$

*Ilo*: $\dfrac{(D_{il})(1-\hat{q}_l)(1-D_{lo})(1-\hat{q}_o)\left(\dfrac{1-D_{lo}C}{1-D_{lo}}\right)}{2}$

***ILo***: $\dfrac{(1-D_{il})(\hat{q}_l)(D_{lo})(1-\hat{q}_o)\left(\dfrac{1-D_{il}C}{1-D_{il}}\right)}{2}$

*ilO*: $\dfrac{(1-D_{il})(1-\hat{q}_l)(D_{lo})(\hat{q}_o)\left(\dfrac{1-D_{il}C}{1-D_{il}}\right)}{2}$

*iLo*: $\dfrac{(D_{il})(\hat{q}_l)(D_{lo})(1-\hat{q}_o)C}{2}$

*IlO*: $\dfrac{(D_{il})(1-\hat{q}_l)(D_{lo})(\hat{q}_o)C}{2}$

B

Given sites A, B, and C, the recombinant fraction between A and B is described by:

$$D_{AB} = D_{AB}(1-D_{BC}) + D_{AB}D_{BC}$$

With interference the consituents are modified by the coefficient of coincidence:

$$D_{AB} = D_{AB}(1-D_{BC})x + D_{AB}D_{BC}C$$

Therefore:

$$x = \frac{1-D_{BC}C}{1-D_{BC}}$$

**Supplementary figure 8.** A. Frequency of allele combinations in three point crosses. The frequency of the double recombinants are modified with coefficient of coincidence (C). The frequency of the single crossovers are reciprocally modified by a factor that is a function of the crossover rate and C. For proof see B.

A

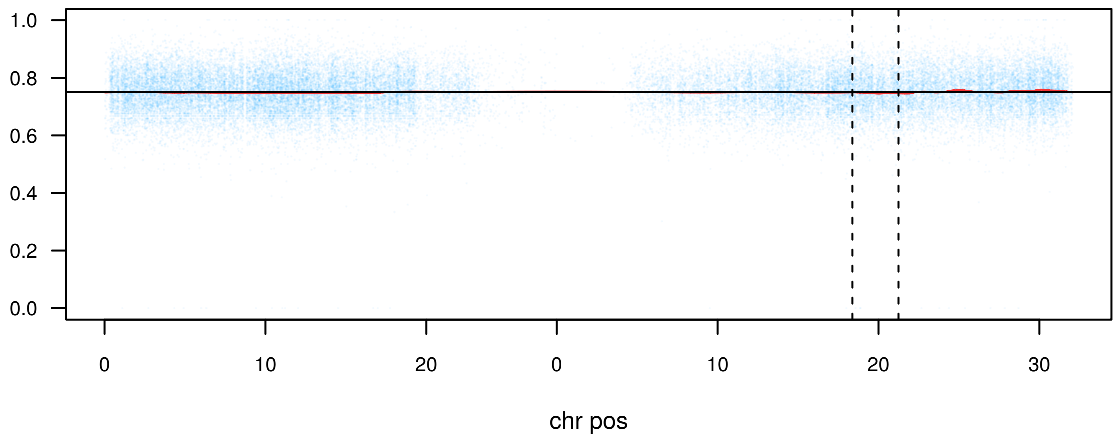


B

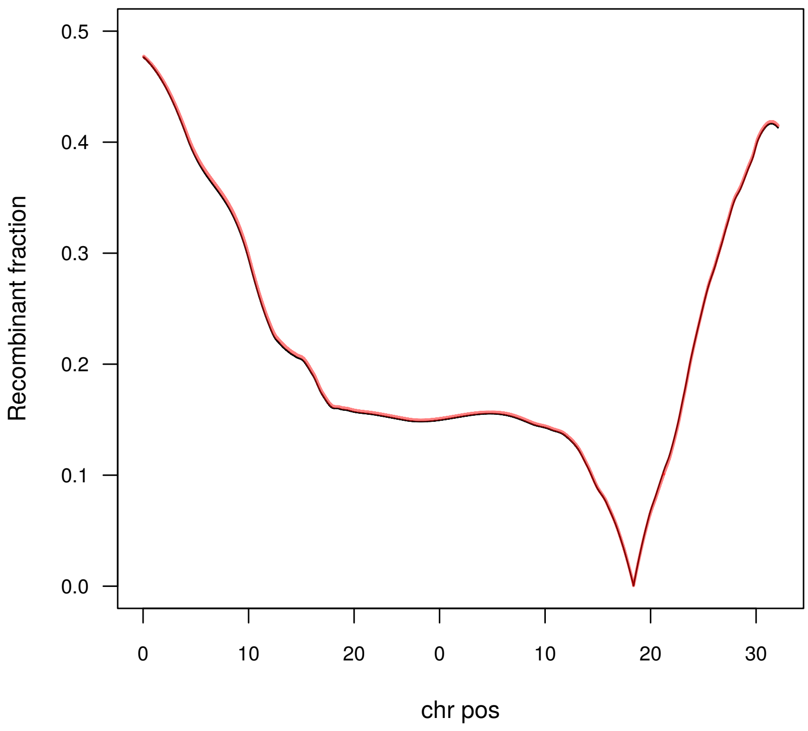


**Supplementary figure 9.** Estimation and correction of offsite viability effect at ebony. A. Viability effect from ebony is determined by summing the allele frequency from both the positive and negative selection pools. The effect equates to -0.015. B. Using this value, we correct the recombinant fraction as according to Figure 4. Black and red lines depict the estimated values before and after correction, respectively.

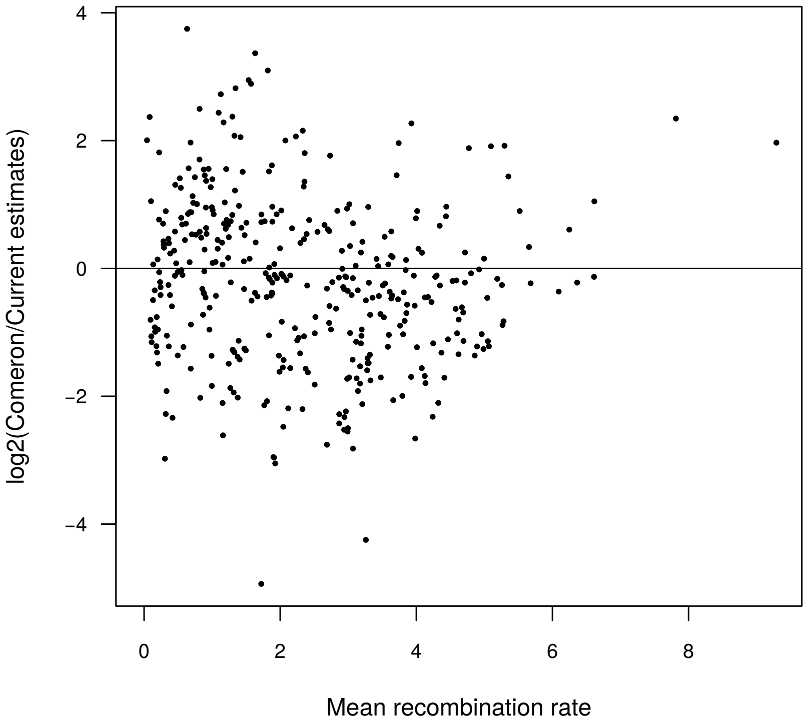


**Supplementary figure 10.** Difference between Comeron et al 2012 estimates and estimates averaged across all marker selected pools. The difference in log scale is plotted on the Y-axis, with the averaged difference between the two methods on the X axis. This is similar to MA-plots for RNA-seq.

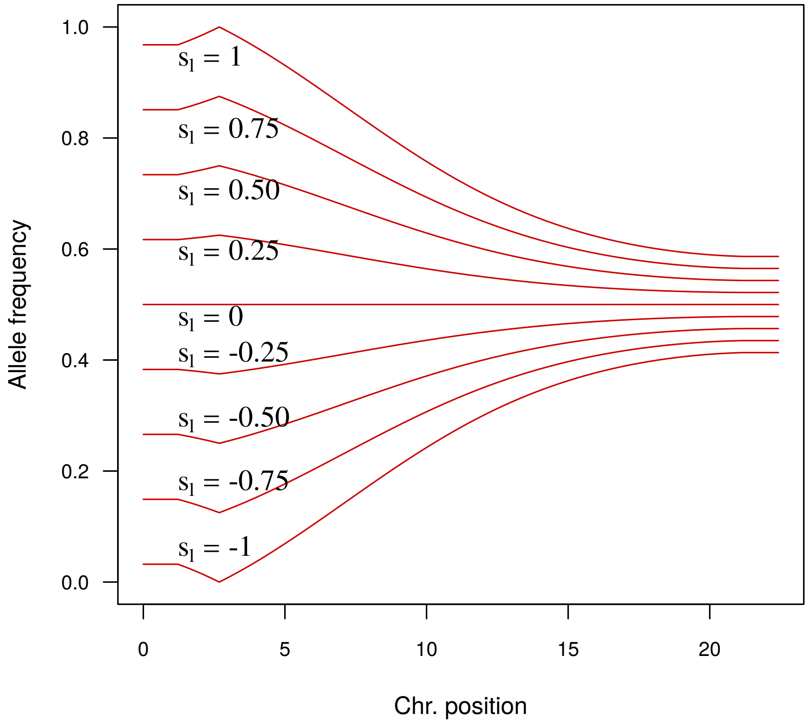


**Supplementary figure 11.** Allele frequency decay at different values of $s_l$ at *w.* Kosambi's mapping function is used to transform genetic distance to recombinant fraction.
